# Separate the wheat from the chaff: genomic scan for local adaptation in the red coral *Corallium rubrum*

**DOI:** 10.1101/306456

**Authors:** M. Pratlong, A. Haguenauer, K. Brener, G. Mitta, E. Toulza, J. Garrabou, N. Bensoussan, P. Pontarotti, D. Aurelle

**Author notes:** These authors jointly supervised this work (P. Pontarotti, D. Aurelle). Corresponding authors /. We have been informed by F. Allendorf that another article had a very similar title: “Waples, R. S. 1998. Separating the wheat from the chaff: Patterns of genetic differentiation in high gene flow species. Journal of Heredity 89:438-450.”.

## Abstract

Genomic data allow an in-depth and renewed study of local adaptation. The red coral (*Corallium rubrum*, Cnidaria) is a highly genetically structured species and a promising model for the study of adaptive processes along an environmental gradient. Here, we used RAD-Sequencing in order to study the vertical genetic structure of this species and to search for signals of local adaptation to depth and thermal regime in the red coral. Previous studies have shown different thermotolerance levels according to depth in this species which could correspond to genetic or environmental differences. We designed a sampling scheme with six pairs of ‘shallow vs deep’ populations distributed in three geographical regions as replicates. Our results showed significant differentiation among locations and among sites separated by around 20 m depth. The tests of association between genetics and environment allowed the identification of candidate loci under selection but with a potentially high rate of false positive. We discuss the methodological obstacles and biases encountered for the detection of selected loci in such a strongly genetically structured species. On this basis, we also discuss the significance of the candidate loci for local adaptation detected in each geographical region and the evolution of red coral populations along environmental gradients.

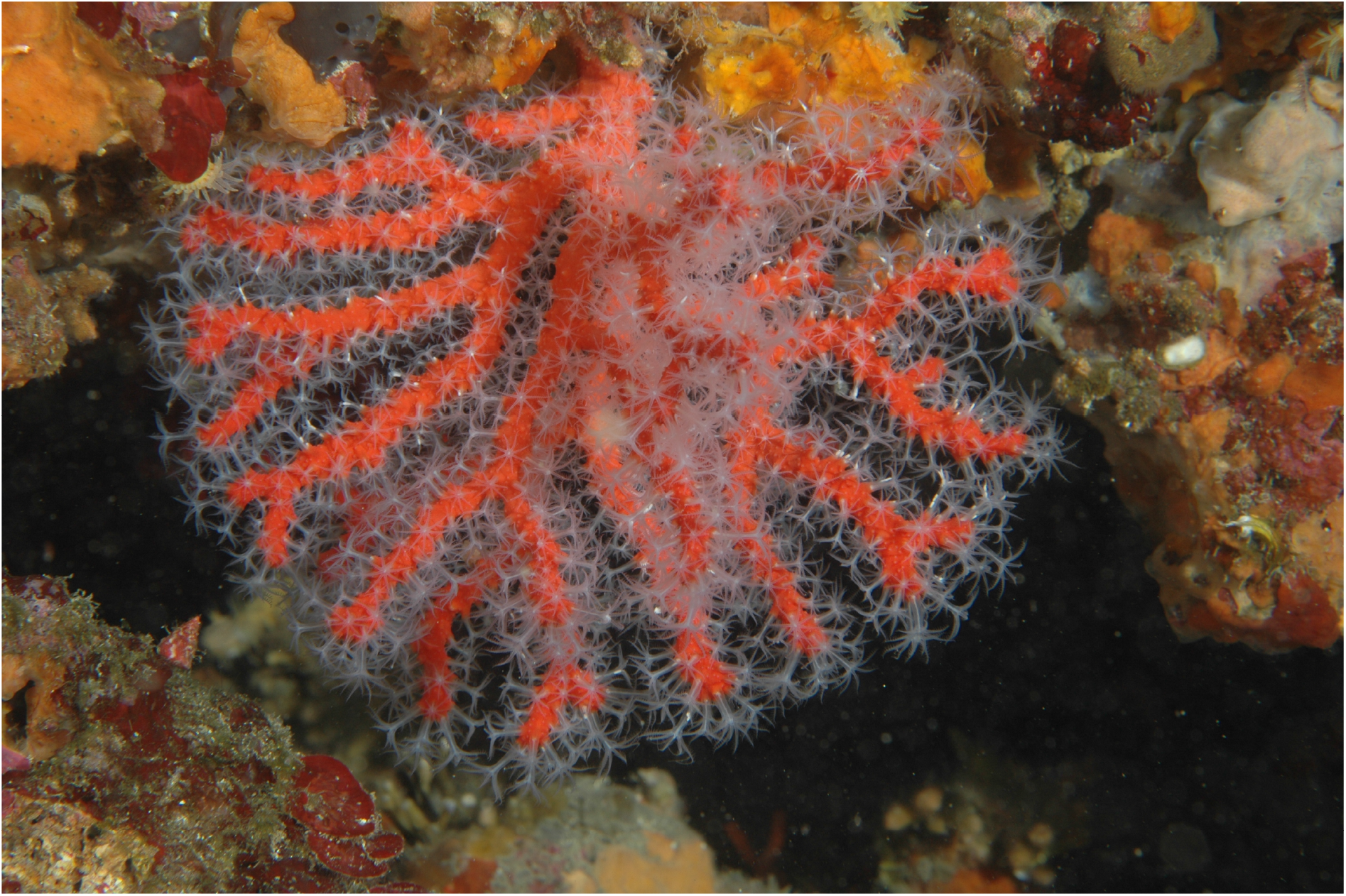
A colony of red coral, *Corallium rubrum*, near Marseille. Photo: F. Zuberer / OSU Pythéas / CNRS

## INTRODUCTION

The study of the mechanisms of adaptation by species to their local environment is of great interest in evolutionary biology. The interaction between environmental conditions, biological traits and evolutionary factors (selection, drift, migration and mutation) will shape the relative importance of genetic and plastic responses for each species facing heterogeneous environmental conditions. If selection is predominant, and if the environmental gradient is persistent for an extended period of time, each local population exposed to local selection could become genetically adapted to the corresponding local environment (Kawecki and Ebert, 2004; Gagnaire and Gaggiotti, 2016). An organism can also cope with local environmental conditions via plasticity or acclimatization, whereby a given genotype develops during its lifetime morphological or physiological responses (DeWitt *et al.*, 1998; Pigliucci, 2001). Although particular situations favoring local adaptation or acclimatization are documented, it is often difficult to disentangle the effects of these two mechanisms and establish their relative contributions to adaptability (Palumbi *et al.*, 2014). In addition, understanding these mechanisms has a fundamental interest in the current context of climate change for improving predictive models and proposing management strategies (Mumby et al., 2011; Gagnaire and Gaggiotti, 2016).

The evolution of adaptive processes depends on the interaction between different processes, including local selection and gene flow. Gene flow can hinder local adaptation through the input in a population of potentially maladapted individuals (migration load; Lenormand, 2002). Conversely, several theoretical studies have shown that gene flow can counteract the effects of genetic drift and promote local adaptation (Hastings and Rohlf, 1974; Felsenstein, 1975; Slatkin and Maruyama, 1975; Nagylaki, 1978; Alleaume-Benharira *et al.*, 2006). The use of high throughput sequencing renewed the study of local adaptation. Various *bottom – up* approaches are now available to study local adaptation through the identification signals of selection along the genome (Barrett and Hoekstra, 2011). In the marine realm, such studies have been conducted at very large scale on highly dispersive teleost species (Bradbury *et al.*, 2010; Limborg *et al.*, 2012; Wang *et al.*, 2013; Milano *et al.*, 2014; Bernardi *et al.*, 2016; Guo *et al.*, 2016), and on benthic invertebrates with a highly dispersive, planctonic larvae stage (Chu *et al.*, 2014; Bay and Palumbi, 2014; Araneda *et al.*, 2016; Benestan *et al.*, 2016). Marine species with high genetic structure are less frequent than more dispersive ones, and genomic studies of local adaptation in such species are still scarce (see Bongaerts *et al.*, 2017 for a recent example). The study of local adaptation in a context of high genetic structure may also be difficult from a methodological point of view: high average F_ST_ values can lead to a high number of false positives in outlier tests for the detection of selection by the corresponding increase in the variance of F_ST_ values (Bierne *et al.*, 2013; Hoban *et al.*, 2016). Furthermore, in a context of high average genomic differentiation, it could be difficult to identify selected loci with a higher differentiation than expected under the neutral model. Finally, if genetic drift is strong, it can generate outlier loci with apparent correlation with an environmental variable outside any selective effect (Kawecki and Ebert, 2004; Hofer *et al.*, 2009; Coop *et al.*, 2010). Therefore the empirical study of local adaptation in such situation remains often challenging and with few empirical data in the marine realm.

Marine coastal environments offer particularly interesting conditions for studies of local adaptation, because of the gradual changes in environmental conditions along coastline at small scale, the more or less gradual vertical changes from shallow to deep water and the patchy distribution of contrasted habitats at different scales (Sanford and Kelly, 2011; Lundgren *et al.*, 2013; Wrange *et al.*, 2014). This interest promoted studies of local adaptation in coastal ecosystems (Ayre, 1995; Ulstrup and Van Oppen, 2003; Smith et al., 2007; Sherman and Ayre, 2008; Barshis *et al.*, 2010; Bongaerts *et al.*, 2011; Barshis *et al.*, 2013; Lundgren *et al.*, 2013; Kersting *et al.*, 2013; Haguenauer *et al.*, 2013; Ziegler *et al.*, 2014; Palumbi *et al.*, 2014; Bay and Palumbi, 2014; Ledoux *et al.*, 2015; Pivotto *et al*., 2015; Jin *et al.*, 2016; Bongaerts *et al.*, 2017). Studying the genetic basis of local adaptation and the connectivity between habitats, could also give some information on the response to climate change (e.g. Bongaerts *et al.*, 2017). In this context genome scans are powerful approaches to explore adaptive processes in natural populations (Manel *et al.*, 2016).

The red coral (*Corallium rubrum*) is an asymbiotic (without *Symbiodinium*) temperate octocoral distributed from 5 to 1016 m depth in the Mediterranean sea and the near Atlantic (Boavida *et al.*, 2016; Knittweis *et al.*, 2016). It is a sessile and long-living species (more than 100 years), with low growth and recruitments rates (Marschal *et al.*, 2004; Santangelo *et al.*, 2012). This species is included in two international conventions (annex III of the Bern Convention and annex V of the European Union Habitats directive), and its harvesting is regulated by national legislations (Ledoux *et al.*, 2013). The study of a few microsatellite loci has demonstrated a strong genetic structure in this species (Ledoux *et al.*, 2010a; Ledoux, *et al.*, 2010b). The shallowest populations, above the seasonal thermocline, are exposed to high maximum temperatures and to frequent and intense thermal fluctuations in summer (Haguenauer *et al.*, 2013). The intensity and frequency of extreme thermal events decrease with depth, and the deepest populations are exposed to stable thermal regimes. Since the observation of mass mortality events affecting this species during thermal anomalies in 1999 and 2003, the thermotolerance of the red coral has been intensively studied in the region of Marseille (France; Garrabou *et al.*, 2001, 2009). Common garden experiments highlighted differences in polyp activities, calcification rate, necrosis rate and expression of HSP70 between shallow and deep individuals (10 or 20 m compared to 40 m depth) facing thermal stress (Torrents *et al.*, 2008; Ledoux *et al.* 2015; Haguenauer *et al.*, 2013). Transcriptomes of individuals from 5 and 40 m were compared and several genes were detected as differentially expressed without the application of any stress (Pratlong *et al.*, 2015). These results suggested the possibility of local adaptation to depth in this species, but the possibility of environmental effects could not be excluded.

Together, these studies highlighted phenotypic differences in thermotolerance levels between individuals from different depths in Marseille, with shallower individuals more tolerant than deeper ones. Nevertheless we still do not know if these differences are the result of local adaptation or of individual acclimatization, or both. Previous works on this species enabled us to have a precise idea of the geographic scale at which local adaptation may occur, and were useful to optimize our sampling design. Because populations from different regions may have evolved similar responses to thermal stress, through similar or different genetic basis, it is interesting to investigate local adaptation in pairs of ‘shallow vs deep’ populations exposed to contrasted thermal regimes in distinct geographical regions (Jones *et al.*, 2012; Hoban *et al.*, 2016). Finally, the study of the genetic structure of this species would be useful to better understand the potential role of deeper populations in reseeding shallower ones following disturbances (Bongaerts *et al.*, 2017).

Here we applied Restriction site Associated DNA sequencing (RAD-Seq) to individuals from pairs of ‘shallow *vs.* deep’ populations in three geographical regions of the Mediterranean Sea. The goal of this study was to characterize the neutral and adaptive genomic variation in this species and to test the possibility of local adaptation to depth through a genome scan approach. Our results enable us to discuss the neutral genetic structure of the red coral. Then we highlight the methodological obstacles expected in the detection of local adaptation in this context. Finally, we discuss the robustness of the candidates of local adaptation detected in each geographical region.

## MATERIAL AND METHODS

### Sampling and DNA extraction

*Corallium rubrum* colonies were collected by scuba diving at two depths of two sites in three geographical regions (Marseille, Banyuls, Corsica) between February and August 2013 (Fig. S1, Table 1). Red coral populations from these three regions correspond to different genetic clusters according to microsatellites (Ledoux *et al.*, 2010b) and RAD-Seq (see results). The two depths of each site presented contrasted thermal regimes with higher mean, maximum and standard deviation of temperature at shallower depths (surveys from March 2012 to October 2014; Table 2). Samples from the two depths at each site will be referred as shallow and deep. The three geographical regions presented different annual variations of temperatures between the two studied depths: a difference of 3.8 °C between the maximum observed at the two depths in Marseille, 1.7 °C in Corsica and 0.5 °C in Banyuls (Table 2). Thirty individuals per site and depth were collected (total 360 individuals), preserved in 95 % ethanol and stored at −20 °C until DNA extraction. Total genomic DNA was extracted according to the protocol of Sambrook *et al.* (1989), followed by a purification using Qiagen DNeasy blood and tissue spin columns (Qiagen). Genomic DNA concentration was quantified using a Qubit 2.0 Fluorometer (Life Technologies).

**Table 1.**
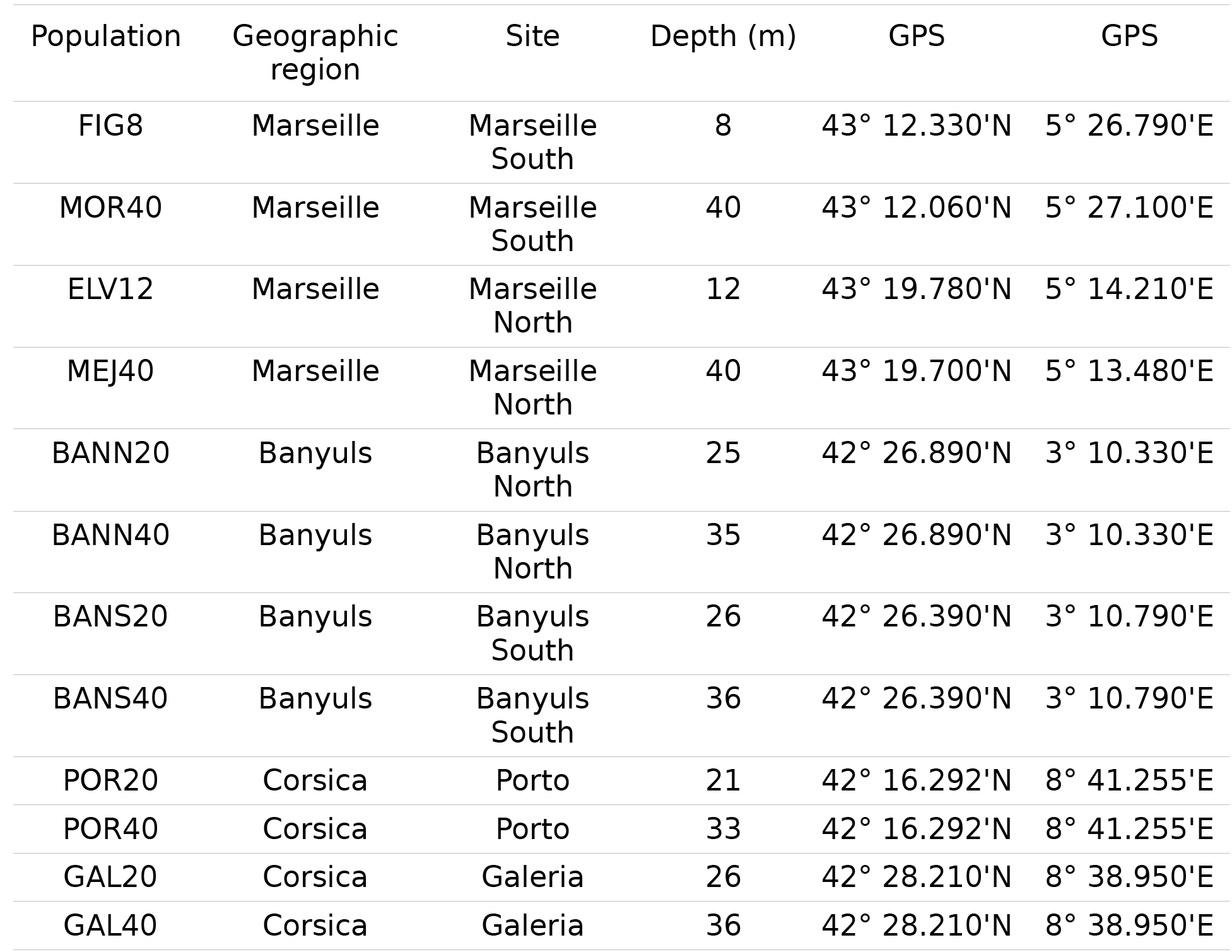
Characteristics of red coral sampling sites.

| Population | Geographic region | Site | Depth (m) | GPS | GPS |
| --- | --- | --- | --- | --- | --- |
| FIG8 | Marseille | Marseille South | 8 | 43° 12.330'N | 5° 26.790'E |
| MOR40 | Marseille | Marseille South | 40 | 43° 12.060'N | 5° 27.100'E |
| ELV12 | Marseille | Marseille North | 12 | 43° 19.780'N | 5° 14.210'E |
| MEJ40 | Marseille | Marseille North | 40 | 43° 19.700'N | 5° 13.480'E |
| BANN20 | Banyuls | Banyuls North | 25 | 42° 26.890'N | 3° 10.330'E |
| BANN40 | Banyuls | Banyuls North | 35 | 42° 26.890'N | 3° 10.330'E |
| BANS20 | Banyuls | Banyuls South | 26 | 42° 26.390'N | 3° 10.790'E |
| BANS40 | Banyuls | Banyuls South | 36 | 42° 26.390'N | 3° 10.790'E |
| POR20 | Corsica | Porto | 21 | 42° 16.292'N | 8° 41.255'E |
| POR40 | Corsica | Porto | 33 | 42° 16.292'N | 8° 41.255'E |
| GAL20 | Corsica | Galeria | 26 | 42° 28.210'N | 8° 38.950'E |
| GAL40 | Corsica | Galeria | 36 | 42° 28.210'N | 8° 38.950'E |

**Table 2.** Temperatures (in °C) characteristics of the sampling sites from March 2012 to October 2014. The corresponding regions are indicated by a letter: M - Marseille, B - Banyuls, C - Corsica

|  | Depth (m) | Minimum | Maximum | Mean | Standard Deviation |
| --- | --- | --- | --- | --- | --- |
| FIG8 (M) | 8 | 12.63 | 26.92 | 17.03 | 3.52 |
| MOR40 (M) | 40 | 12.73 | 23.06 | 15.40 | 2.11 |
| ELV12 (M) | 12 | 11.81 | 26.70 | 16.60 | 3.24 |
| MEJ40 (M) | 40 | 11.86 | 22.87 | 15.29 | 2.18 |
| BANN20 (B) | 25 | 12.22 | 24.29 | 17.20 | 2.63 |
| BANN40 (B) | 35 | 9.41 | 23.83 | 14.49 | 2.45 |
| BANS20 (B) | 26 | 12.22 | 24.29 | 17.20 | 2.63 |
| BANS40 (B) | 36 | 9.41 | 23.83 | 14.49 | 2.45 |
| POR20 (C) | 21 | 12.51 | 25.91 | 17.51 | 3.41 |
| POR40 (C) | 33 | 12.56 | 23.83 | 16.26 | 2.45 |
| GAL20 (C) | 26 | 12.46 | 25.09 | 17.13 | 3.13 |
| GAL40 (C) | 36 | 12.56 | 23.83 | 16.26 | 2.45 |

### RAD-Sequencing

Twelve RAD libraries were prepared according to the protocol described in Etter *et al.* (2011), with small modifications. Briefly, 1 μg of genomic DNA for each sample was digested using high-fidelity PstI during 60 min at 37 °C. P1 adapters, with 4-6 bp individual barcodes were then ligated to each sample using 0.5 μL of T4 DNA ligase (NEB), 0.5 μL of rATP 100 mM (Promega), 1 μL of DTT 500 mM (Promega), 1 μL of 10X T4 ligase buffer (NEB) and incubated during 60 min at 22 °C, 10 min at 65 °C and 1 min at 64 °C. Individual samples were pooled by 32 (generally by location), sheared, size selected and P2-barcoded. Final PCR for RAD-tags enrichment were performed with 16 cycles and primers dimers were removed during a final AMPure Beads Purification (Agencourt). Libraries were sequenced on an Illumina HiSeq2000 using 100 bp single-end reads, at the Biology Institute of Lille (IBL, UMR 8199 CNRS) and at the MGX sequencing platform in Montpellier (France).

The STACKS pipeline (Catchen *et al.*, 2011, 2013) was used for the loci *de novo* assembly and genotyping. Quality filtering and demultiplexing were performed with the *process_radtags* module with default parameters which enables to remove any read with uncalled base and to perform a phred-33 quality filtering of raw reads. Exact-matching RAD loci (putative orthologous tags) were individually assembled using *ustacks* with a minimum depth of coverage of five reads per allele (m = 5) and a maximum of five nucleotide mismatches between allele (M = 5). These parameters were optimized during preliminary runs. *Cstacks* was used to build a catalog of consensus loci from all individuals, with five mismatches allowed between individuals at the same locus (n = 5). Matches of individual RAD loci to the catalog of loci were searched using *sstacks*. Finally, the *population* module was used to obtain the loci that were successfully genotyped in at least 75 % of individuals from all populations. We observed an increase in the number of SNPs from position 86 bp to 91 bp and we removed these positions from the analysis which were due to sequencing problems. In order to filter for poor-quality SNPs and artifacts due paralogous sequences, we used VCFtools (Danecek *et al.*, 2011) to remove SNPs that were not at the Hardy-Weinberg equilibrium within at least one of the 12 populations with a p-value threshold of 0.01. SNPs with a minor allele frequencies below 0.01 were removed using VCFTools. Individuals with more than 30 % of missing genotypes were discarded. Finally, only the first SNP of each RAD locus was kept for further analysis. The whole dataset has been previously used for the study of sex determinism in *C. rubrum* (Pratlong *et al.*, 2017); we develop here the study of genetic structure and local adaptation.

### Diversity and neutral genetic structure

Global F_IS_ over alleles and expected heterozygosity were estimated using GENEPOP and ARLEQUIN v.3.5 (Rousset, 2008; Excoffier and Lischer, 2010). The *C. rubrum* genetic structure was first analyzed by principal component analysis (PCA) using the package adegenet in R (Jombart, 2008; R Core Team, 2016). This analysis was performed on the total dataset (12 populations) and inside each of the three studied geographical regions (four populations in each). The dataset was centered and missing data were replaced by the mean allele frequency for each locus (http://adegenet.r-forge.r-project.org/files/tutorial-basics.pdf). In a second step, we performed a Bayesian population clustering implemented in the program STRUCTURE v.2.3.4 (Pritchard *et al.*, 2000; Falush *et al.*, 2003, 2007; Hubisz *et al.*, 2009). We performed ten independent replicates from *K* = 1 to 10 with a burn-in of 50 000 and a number of MCMC iterations after burn-in of 100 000, with the model allowing for admixture and correlated allele frequencies. We calculated the Δ*K* statistic of Evanno *et al.* (2005) to help in the choice of the most appropriate number of genetic clusters but we also considered different K values. We used CLUMPAK to summarize the STRUCTURE results from the ten independent runs (Kopelman *et al.*, 2015). The global and pairwise populations F_ST_ and exact tests for population differentiation were computed with GENEPOP 4.0.10 (Rousset, 2008). The correlation between the spatial distance between the two depths of the same site and the corresponding population pairwise F_ST_ was tested with the correlation test of Spearman implemented in R (R Core Team, 2016). Finally, we conducted an analysis of molecular variance (AMOVA) in ARLEQUIN v.3.5 (Excoffier and Lischer, 2010) with 10 000 permutations. The hierarchy for this analysis was chosen to follow the three geographical regions of our samples (Marseille, Corsica and Banyuls). This choice was justified by the PCA on the overall dataset. Finally, we performed the PCA and F_ST_ calculation using a dataset comprising only putatively neutral SNPs (without the SNPs detected as outliers by F_ST_ outlier methods, see below).

### Detection of local adaptation

In order to search for loci potentially involved in local adaptation, we first used BayeScEnv (Villemereuil and Gaggiotti, 2015). This method identifies F_ST_ outlier loci that show a relationship between genetic differentiation and environmental differentiation. Runs were performed using default parameters, except the number of pilot runs that was set at 40. The maximal temperature recorded in each site was used as environmental variable (Table 2). We tested other descriptors of the thermal regime and we got similar results (data not shown). The convergence of runs was checked with the Gelman and Rubin’s diagnostic using the R package coda (Plummer *et al.*, 2006).

Second, we searched for F_ST_ outliers among red coral populations using ARLEQUIN v.3.5 (Hofer *et al.*, 2009; Excoffier and Lischer, 2010). Because hierarchical genetic structures are known to lead to a high number of false positives in the search of outlier loci (Hofer *et al.*, 2009), we performed this analysis independently in the three geographical regions in order to down a level in the structure. With this method, a distribution of F_ST_ across loci as a function of heterozygosity between populations is obtained by performing simulations under a hierarchical island model (two depths in one site and two sites in one geographical region). Outliers were identified as loci being in the tails of the generated distribution (p < 0.01). Outliers detected by ARLEQUIN could be false positives or the result of a selective pressure independent of depth. Therefore, we selected among these candidate loci, those linked with depth differentiation by searching, inside each geographical regions, loci with significant differences in genotypic frequencies between depths according to a Chi² test. Significant differences were identified by using a false discovery rate of 0.05 (Benjamini and Hochberg, 1995).

Finally, we used the R package pcadapt to search for outliers loci by taking into account population structure and individual admixture (Luu *et al.*, 2017). This method is recommended in cases of hierarchical genetic structure for a better control of the false positive rate. By identifying outliers loci linked with a particular principal component, pcadapt enabled us to focus on candidates linked with our biological question. From the pcadapt analyses, we selected outliers candidates linked with the relevant principal components with a q-value cutoff of 0.01.

### Functional annotation and enrichment tests

The RAD tags were aligned on the red coral transcriptome (Pratlong *et al.*, 2015) using the Burrows-Wheeler Alignment Tool (BWA; Li and Durbin, 2009). Blast2GO was used for the annotation of resulting contigs and functional enrichment analysis (Conesa *et al.*, 2005). First, a blastp was first performed on the NCBI nr database with an e-value threshold of 10^-10^ (Altschul *et al.*, 1990). Then, Blast2GO retrieved Gene Ontology (GO) terms associated with the obtained BLAST hits. Finally, in order to identify function potentially over-represented in outliers, we performed an enrichment analysis using a Fisher’s exact test corrected using a false discovery rate of 0.05 (Benjamini and Hochberg, 1995).

## RESULTS

### RAD-Sequencing and genotyping

An average of 191 ± 21 millions of reads by library was obtained after sequencing. After the demultiplexing and cleaning processes of the STACKS’s *process_radtags* module, an average of 180 ± 22 millions of reads by library was obtained. From these reads, we were able to assemble 138 810 unique consensus RAD-tags present in at least 75 % of our 360 individuals. After all quality filter steps (Table 3), 27 461 SNPs were available. The mean proportion of missing data per locus before filtering for MAF 0.01 was 4.22 % (standard error: 2.78%). Finally, we removed six individuals presenting more than 30 % of missing data (one individual from the MEJ40 population, two from the BANN40 population, two from the GAL20 population and one from the GAL40 population). Our final dataset used for further analysis consisted in 359 individuals genotyped on 27 461 SNPs.

**Table 3.** Counts of SNP loci after each filtering step.

| Step | Number of SNPs | Software |
| --- | --- | --- |
| After assembly raw data | 138 810 | Stacks<br>(Catchen *et al.*, 2011, 2013) |
| Excluding loci not in<br>within population HWE | 86 520 | VCFtools<br>(Danecek *et al.*, 2011) |
| MAF 1 % | 56 844 | VCFtools<br>(Danecek *et al.*, 2011) |
| One SNPs per RAD-tag | 27 461 |  |

### Genetic diversity

Multilocus values of the F_IS_ ranged between 0.005 (ELV12) and 0.065 (BANS40) (Table 4). Expected heterozygosity varied from 0.12 (GAL20) to 0.18 (all populations of Marseille) (Table 4). Populations of Marseille had higher values of expected heterozygosity than populations from Corsica and Banyuls (p = 0.02, Wilcoxon–Mann–Whitney test).

**Table 4.** Summary statistics of genetic diversity. The nucleotide diversities and numbers of private alleles are based on the dataset with all SNPs for each RAD-loci, excluding those with the lowest quality, and the removal of loci with minor allele frequencies below 0.01 (total 56 844 SNPs). Measures of F_IS_ and expected heterozygosities were based on the final dataset comprising 27 461 SNPs (with one SNP / locus). The corresponding regions are indicated by a letter: M – Marseille, B – Banyuls, C – Corsica

| Population | Sample size | Nucleotide diversity | Average number of private alleles | $F_{IS}$ | Expected heterozygosity |
| --- | --- | --- | --- | --- | --- |
| BANN20 (B) | 30 | 0.172 | 0 | 0.018 | 0.15 |
| BANN40 (B) | 28 | 0.174 | 0 | 0.012 | 0.15 |
| BANS20 (B) | 30 | 0.173 | 0 | 0.019 | 0.15 |
| BANS40 (B) | 30 | 0.172 | 0 | 0.065 | 0.13 |
| ELV12 (M) | 30 | 0.192 | 0.001 | 0.005 | 0.17 |
| MEJ40 (M) | 29 | 0.195 | 0 | 0.053 | 0.18 |
| FIG8 (M) | 30 | 0.192 | 0.001 | 0.005 | 0.18 |
| MOR40 (M) | 30 | 0.198 | 0.001 | 0.036 | 0.18 |
| GAL20 (C) | 28 | 0.124 | 0.002 | 0.013 | 0.09 |
| GAL40 (C) | 29 | 0.149 | 0.001 | 0.019 | 0.13 |
| POR20 (C) | 30 | 0.165 | 0 | 0.023 | 0.13 |
| POR40 (C) | 30 | 0.168 | 0.006 | 0.009 | 0.15 |

### Population structure analysis

The positioning of individuals with respect to the first two principal components reflected the geographical, and, partly, the depth origin of the individuals (Fig. 1A). Individuals from Marseille and from Banyuls formed two clear and homogeneous groups while individuals from the two sites of Corsica formed two different groups with an important distance between them on the second axis. The first PCA axis explained 7.28 % of the total genotypic variance and separated individuals from Marseille from individuals from Banyuls and Corsica. The second axis explained 4.4 % of the total genetic variance and separated individuals from the Porto site in Corsica from other individuals. The fifth axis of the PCA separated all individuals according to their sex, independently from their geographical origin, as observed in a previous study (Pratlong et al., 2017). This axis can be explained by loci diagnostic of sex, heterozygous in females and homozygous in males (see Pratlong et al., 2017). Concerning PCA inside geographical regions, individuals from the two sites of Corsica and Marseille (north and south) were separated along the first axis (13.41 % and 6.74 % of the total genetic variance respectively; Fig. 1B and 1C). The second axis (2.77 % of the total genetic variance) separated populations from the two depths of the two sites of Marseille. In Corsica and Banyuls, no PCA axis showed clear association with depth. Individuals from the two depths of the Galeria (GAL) site of Corsica were separated along the second axis (4.02 % of the total genetic variance) but this was not the case for individuals from the two depths of the Porto site (POR). Individuals from Banyuls showed much less structure than individuals from Marseille and Corsica (Fig. 1D). The first axis (2.99 % of the total genetic variance) separated individuals from the two sites (north and south). The second axis (2.46 % of the total genetic variance) separated individuals according to their sex (Pratlong *et al.*, 2017). The PCA on the overall dataset and inside each geographical region gave similar results when only putatively neutral SNPs were considered (Fig. S2).

**Figure 1.**
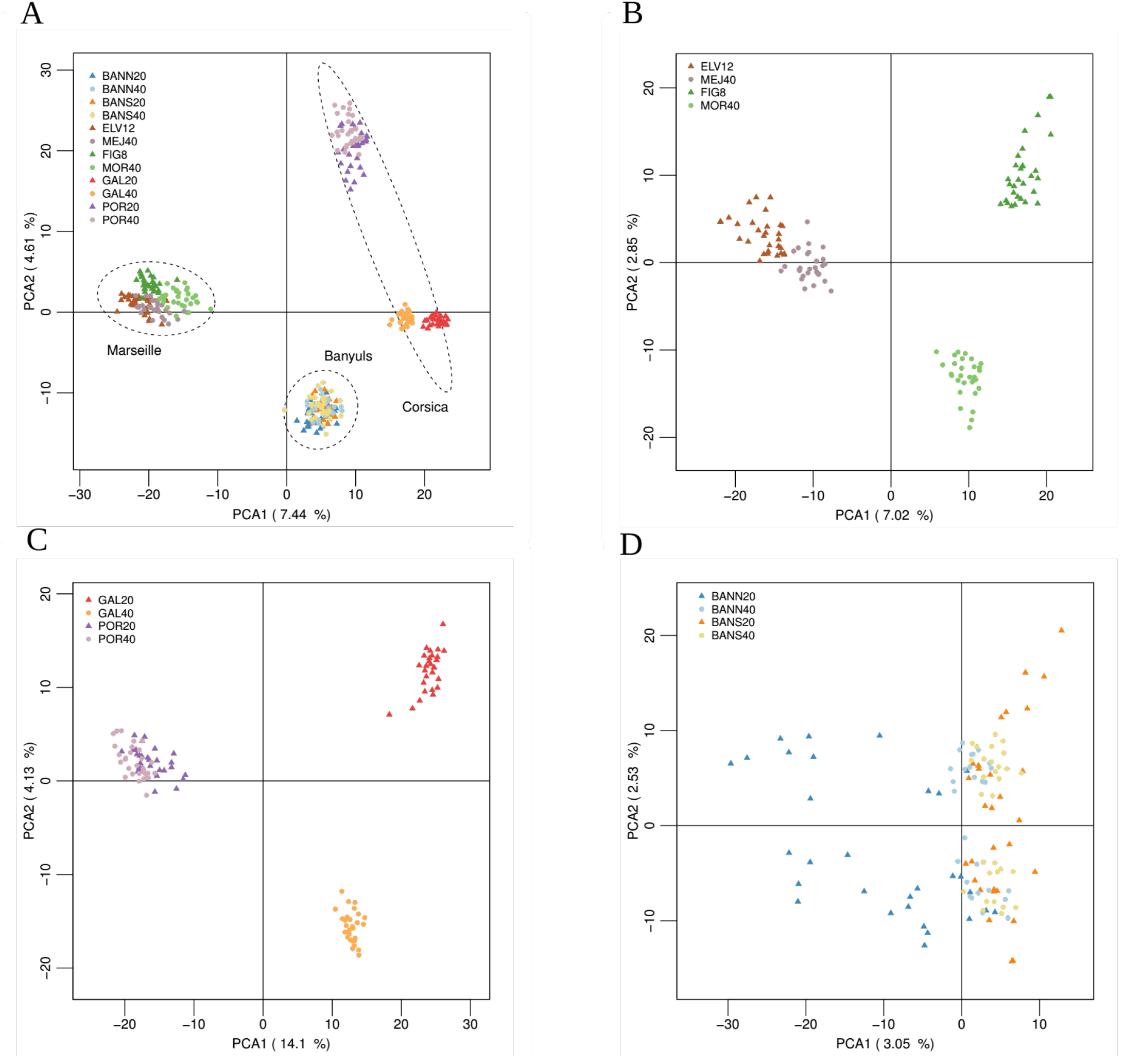
Principal component analysis (Axes 1 and 2) of A) the 12 red coral populations (n = 354 individuals, 27 461 SNPs), B) the four red coral populations from Marseille (n =119 individuals, 27 461 SNPs), C) the four red coral populations from Corsica (n =117 individuals, 27 461 SNPs), D) the four red coral populations from Banyuls (n =118 individuals, 27 461 SNPs).

The delta(K) criterion (Evanno *et al.*, 2005) indicated K = 2 as the most informative number of clusters for the STRUCTURE analysis. We present here the results for K =2 to K =4 which captured the main information (Fig. 2). In all cases, all clusters corresponded to the main geographical boundaries and the two depths of each site always clustered together. For K = 2, a clear separation between the Marseille regions and the Corsica / Banyuls regions was observed, confirming the separation of populations along the first PCA axis (Fig. 1). The clustering at K = 3 separated the three geographical regions in 7/10 replicates, and the remaining replicates grouped either one or the other Corsican sites with Banyuls populations (Fig. S3). Finally, K = 4 separated the two Corsican sites.

**Figure 2.**
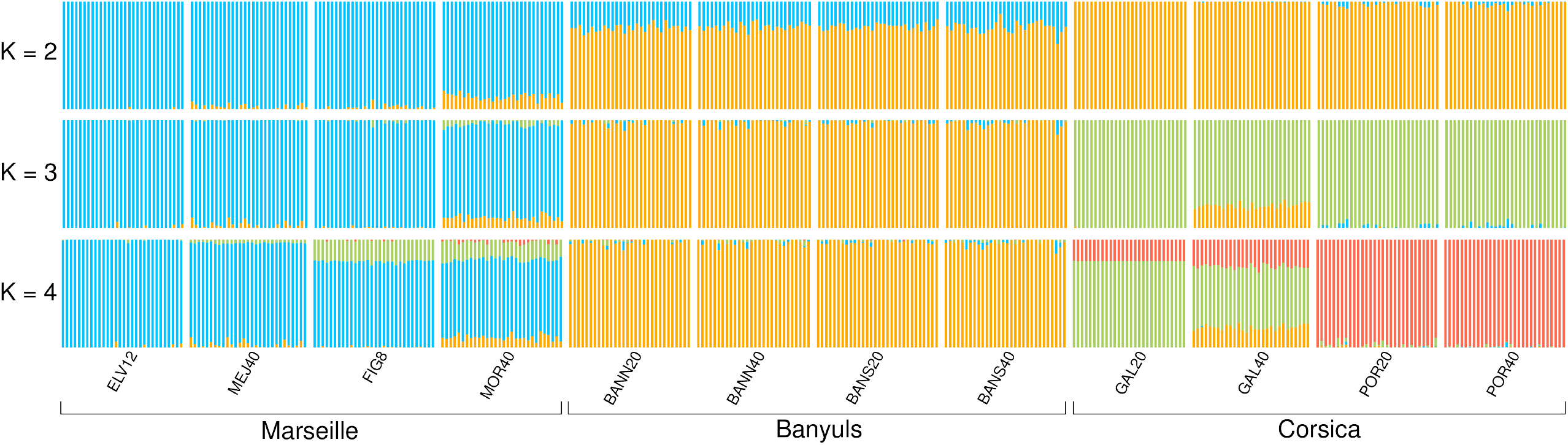
Results from Bayesian individual clustering with STRUCTURE for K = 2 to K = 4. For K = 2 and K = 4, all ten replicates produced the same structure. For K = 3, the major mode presented here was the result of 7/10 replicates. Minor modes are presented in Fig. S3.

The overall multilocus F_ST_ of the total dataset was 0.13. Pairwise F_ST_ values ranged from 0.01 (BANS20 vs BANS40, BANS20 vs BANN40 and BANN40 vs BANS40) to 0.24 (ELV12 vs GAL20 and FIG8 vs GAL20; Table 5). The exact test of genetic differentiation was highly significant for all pairwise comparisons (p < 0.001), even for the Banyuls and Galeria populations separated by 10 m depth (F_ST_ = 0.022 for BANN20 vs BANN40, F_ST_ = 0.012 for BANS20 vs BANS40 and F_ST_ = 0.10 for GAL20 vs GAL40). Considering the F_ST_ between depths, high F_ST_ values can be observed for different loci and different samples comparisons (Fig. S4). The average F_ST_ between the two depths of the same site was 0.04 in Marseille, 0.08 in Corsica and 0.02 in Banyuls (0.04 for the total dataset). Considering only putatively neutral loci (see below for outliers loci), the overall F_ST_ of the total dataset was 0.12 and pairwise F_ST_ values ranged from 0.01 (BANS20 vs BANS40, BANS20 vs BANN40 and BANN40 vs BANS40) to 0.23 (ELV12 vs GAL20) (Table S1). There was no correlation between the distance between two depths of the same site and the corresponding population pairwise F_ST_ (p = 0.75). We obtained a similar result (p = 1) if we removed the four populations of Marseille whose sampling sites for the two considered depths were not exactly the same (653 m horizontal distance between FIG8 and MOR40 and 995 m between ELV12 and MEJ40). Finally, the F_ST_ between the two shallow sites inside a geographical region was in all three cases higher than those between the two deep sites of the same region (0.10 vs 0.058 in Marseille, 0.20 vs 0.14 in Corsica and 0.025 vs 0.014 in Banyuls. This difference was significant in each of the three regions (p < 1.10^-16^ with a t-test in all cases). In a similar way, the F_ST_ between two shallow sites of two different geographical regions were in all cases higher than those between the two corresponding deep sites, except for the comparisons between the Porto sites and the Banyuls sites (p = 0.38 and p = 0.02 for the POR/BANN and POR/BANS comparisons respectively; p < 1.10^-16^ for the other comparisons).

**Table 5.**
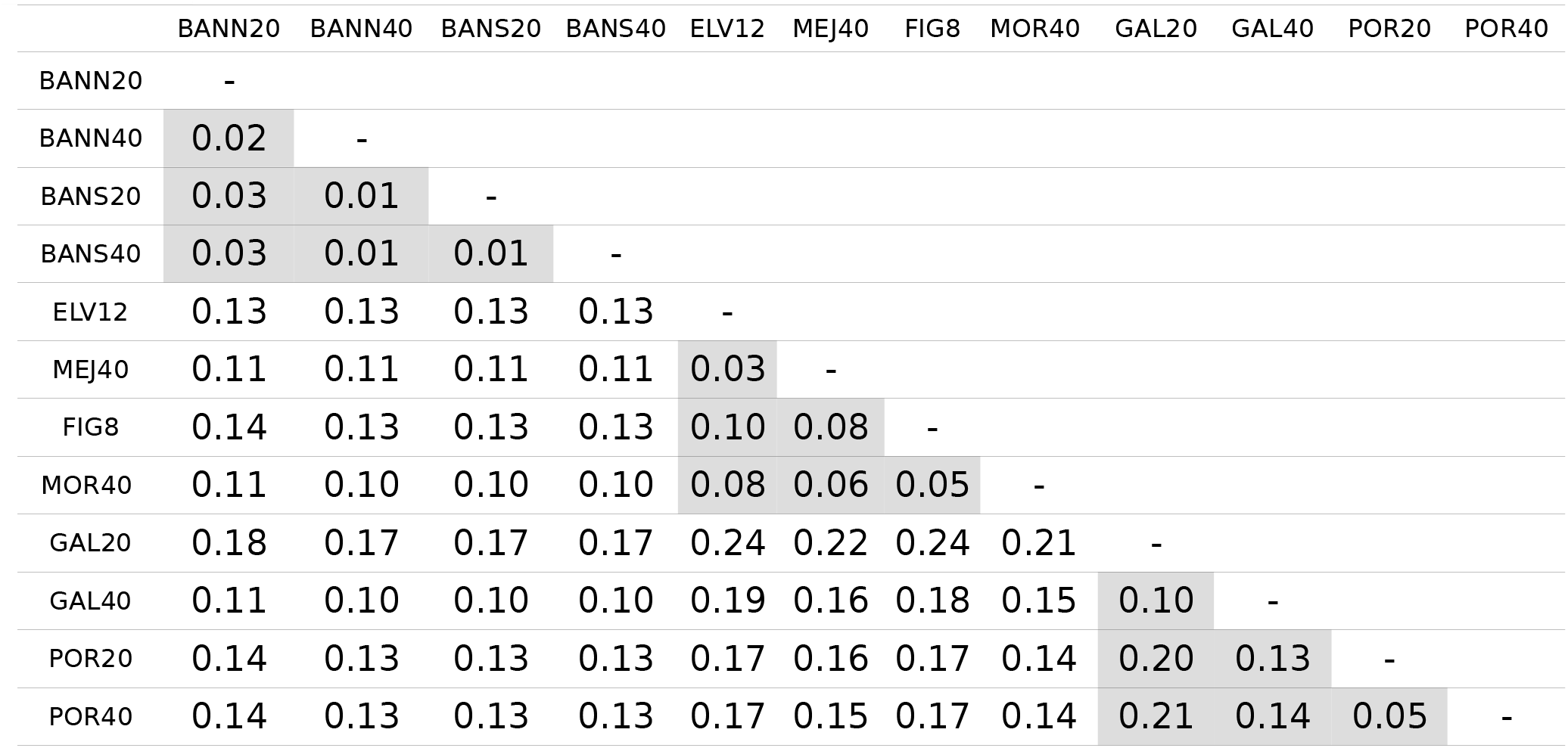
Pairwise F_ST_ estimates. All comparisons were highly significant. Intra-region comparisons are highlighted.

|  | BANN20 | BANN40 | BANS20 | BANS40 | ELV12 | MEJ40 | FIG8 | MOR40 | GAL20 | GAL40 | POR20 | POR40 |
| --- | --- | --- | --- | --- | --- | --- | --- | --- | --- | --- | --- | --- |
| BANN20 | - |  |  |  |  |  |  |  |  |  |  |  |
| BANN40 | 0.02 | - |  |  |  |  |  |  |  |  |  |  |
| BANS20 | 0.03 | 0.01 | - |  |  |  |  |  |  |  |  |  |
| BANS40 | 0.03 | 0.01 | 0.01 | - |  |  |  |  |  |  |  |  |
| ELV12 | 0.13 | 0.13 | 0.13 | 0.13 | - |  |  |  |  |  |  |  |
| MEJ40 | 0.11 | 0.11 | 0.11 | 0.11 | 0.03 | - |  |  |  |  |  |  |
| FIG8 | 0.14 | 0.13 | 0.13 | 0.13 | 0.10 | 0.08 | - |  |  |  |  |  |
| MOR40 | 0.11 | 0.10 | 0.10 | 0.10 | 0.08 | 0.06 | 0.05 | - |  |  |  |  |
| GAL20 | 0.18 | 0.17 | 0.17 | 0.17 | 0.24 | 0.22 | 0.24 | 0.21 | - |  |  |  |
| GAL40 | 0.11 | 0.10 | 0.10 | 0.10 | 0.19 | 0.16 | 0.18 | 0.15 | 0.10 | - |  |  |
| POR20 | 0.14 | 0.13 | 0.13 | 0.13 | 0.17 | 0.16 | 0.17 | 0.14 | 0.20 | 0.13 | - |  |
| POR40 | 0.14 | 0.13 | 0.13 | 0.13 | 0.17 | 0.15 | 0.17 | 0.14 | 0.21 | 0.14 | 0.05 | - |

The AMOVA indicated a similar percentage of the molecular variance explicated by differences among group and within groups (7.8 and 7.07 % respectively) and approximately 85 % of variance explicated by differences within populations (Table 6). There was significant genetic differentiation at the three studied levels (F_ST_ = 0.15, F_SC_ = 0.08, F_CT_ = 0.08; p < 0.001 in the three cases).

**Table 6.** Percent of the variation explained by grouping populations according to their geographical region on the analysis of molecular variance (performed with ARLEQUIN).

| Source of variation | d.f. | Percentage of variation |
| --- | --- | --- |
| Among groups | 2 | 7.80 |
| Among populations within groups | 9 | 7.07 |
| Within populations | 696 | 85.13 |

### Outliers SNPs

We identified 82 outliers with BayeScEnv. However, we noticed that all these outliers seemed to be driven by the divergence between particular populations, with one allele being always fixed in one or several populations without logical association with depth. ARLEQUIN detected 563 loci potentially under selection in Marseille, 869 in Corsica and 397 in Banyuls. Among these SNPs, all corresponded to a signal of divergent selection in Marseille and Banyuls, 207 of the 869 candidate loci corresponded to a signal of balanced selection in Corsica and the remaining 662 loci corresponded to a signal of divergent selection. Considering only these outliers, the overall F_ST_ of the total dataset was 0.25 and pairwise F_ST_ values ranged from 0.02 (BANS20 vs BANS40) to 0.42 (GAL20 vs POR40) (Table S2). The 207 loci potentially under balanced selection in Corsica were linked with sex differentiation and were not further analyzed here (Pratlong *et al*. 2017). Eight outlier SNPs were detected both in Marseille and Banyuls, 12 both in Marseille and Corsica and 12 both in Corsica and Banyuls. No SNP was detected as potentially under divergent selection and common in the three regions. The complementary Chi² test of homogeneity of genotypic frequencies between depths inside each region detected 162 candidate loci in Marseille, 1 371 in Corsica and 3 in Banyuls. Among these loci, 35, 248 and 2 where also respectively detected with the ARLEQUIN analysis. The numbers of outlier loci were correlated with the variance and the average of F_ST_ values inside each geographical regions (correlation coefficient of 0.97). The second axis of the PCA using the Marseille individuals showed apparent association with depth and appeared to be influenced by the variation of the candidates for local adaptation to depth detected by ARLEQUIN: 51 % of these loci were in the top 1 % of the contributions to this second axis, and 86 % were in the top 5 %.

Because the Marseille region is the only one presenting a principal component linked with depth (Fig. 1C), the pcadapt results obtained for the Corsica et Banyuls regions have poor biological relevance for our biological question. We chose thus to present only the pcadapt results obtained for the Marseille region. Pcadapt detected 58 outliers loci linked with the second PCA axis, the one which was linked to depth differentiation. All these candidates were detected by the ARLEQUIN analyses and 20 were also common with the Chi² test presented above.

### Functional annotation

Among the 27 461 analyzed RAD-tags, 6 376 had hits on the red coral transcriptome (23.2 %). Concerning SNPs detected as outliers by ARLEQUIN and contributing to the depth divergence, 8 on the 35 detected in Marseille had hits on the transcriptome, 46 on the 248 detected in Corsica and 2 on the 2 detected in Banyuls (Table S3). We did not observed any GO term enriched in coding regions among candidates SNPs, nor any functional enrichment in these outliers.

## DISCUSSION

### Genetic diversity and structure

Our results confirm at a genomic scale the high genetic structure of the red coral, which was observed with a small number of microsatellite loci (Costantini *et al.*, 2007; Ledoux *et al.*, 2010b). Here the differentiation was significant for all comparisons, even with low average F_ST_ in some cases (0.01 or 0.02 between depths in Banyuls). Such low but significant F_ST_ can be explained by the high number of loci used here, and by the high variance in F_ST_ values. Indeed for all comparisons some important F_ST_ were observed in the dataset (see below). This significant differentiation at all distances could be the consequence of reduced mean larval dispersal distance, despite a quite long pelagic larval duration estimated in aquarium (from 16 to 42 days; Martínez-Quintana *et al.*, 2015). Genetic incompatibilities could also contribute to the observed differentiation at least for some loci (Kulmuni and Westram, 2017). Our analysis of the genetic structure of the red coral revealed several clusters mainly corresponding to the geographical distributions of this species. A high differentiation was observed here with PCA and STRUCTURE between the two sites of Corsica separated by around 22 km. This pattern of genetic structure could be explained by a putative barrier to gene flow (through currents or lack of suitable habitats) between the two Corsican sites, or it could also correspond to an historical separation of these populations: two lineages could then be present in Corsica, with one being related to Banyuls populations.

We reported here a significant vertical genetic structure between the two depths of the same site (populations separated by less than 20 m). This differentiation was also observed when outlier loci were removed indicating that it is also shaped by neutral processes (migration / drift) as suggested previously with microsatellites (e.g. Ledoux *et al.*, 2010a). In a study of the vertical genetic structure of red coral in two western Mediterranean sites (Cap de Creus, Spain and Portofino, Italia), Costantini *et al.* (2011) observed a drop in connectivity around 40 – 50 m depth, with genetic diversity declining with depth, but our sampling scheme did not allow us to test this hypotheses. The study of the vertical genetic structure in corals is important in the context of climate change. As deeper populations may be less affected by climate change, they could possibly reseed shallower populations (Bongaerts *et al.*, 2017). Nevertheless this possibility of reseeding depends on the connectivity or potential barriers between depths. A lack of connectivity could erroneously be inferred in cases of cryptic species (Pante *et al.*, 2015). Contrary to Prada *et al.* (2008) who showed the existence of two cryptic lineages at two different depths in a tropical octocoral, the populations of red coral from the two studied depths clearly correspond here to the same species, and the differentiation between depth was lower than the differentiation between sites and regions. This vertical structure may be the result of both inherent life history traits and environmental variables. Weinberg (1979) has described a negative geotropism for the planulae of *C. rubrum*, and Martínez-Quintana *et al.* (2015) demonstrated that this was an active behavior. Depending on the orientation of the substrate, this can limit the connectivity between depths. The seasonal stratification during larval emission (which can occur from June to September; Haguenauer A., pers. comm.) could also limit dispersal. According to these hypotheses the larval behavior and oceanographic factors should lead to the genome-wide neutral differentiation between depths. This differentiation is probably also shaped by drift induced by the small effective size of red coral populations (Ledoux et al., 2010a).

The horizontal genetic differentiation between the two shallow sites was higher than those between the two corresponding deep sites, inside and between geographical region. This suggests a higher connectivity or lower rate of genetic drift for deep populations compared to shallow ones. The repeated colonization of shallow depths from deeper ones or the higher harvesting pressure on shallow populations compared to deep one could enhance genetic drift as well (Rossi *et al.*, 2008; Cannas *et al.*, 2016). We did not observe here a reduction in expected heterozygosity for shallow populations, connectivity differences then seem to be more probable in explaining the observed differences of genetic differentiation. Interestingly, Rossi *et al.* (2008) observed a higher frequency of patches of red coral below 50 m compared to above 50 m: if such pattern is present in the area and depths considered here, then it could increase gene flow through stepping stones migration. The observed vertical and horizontal genetic structure could indicate reduced recolonization abilities following disturbances such as mortality event induced by heat waves. Nevertheless the observed genetic structure could also be shaped by colonization history and monopolization effect (Orsini *et al.*, 2013). In this case, a disturbance leading to free habitats would facilitate recolonization from other populations.

### Potential biases in the search of outlier loci

We observed high F_ST_ values for different loci and different sample comparisons, and not only between depths (Fig. S4). The methods used here to identify selected loci will most likely detect loci with strong effects (Pritchard and Di Rienzo, 2010; Gagnaire and Gaggiotti, 2016), and it is highly improbable to observe such a high number of selected loci. Both hierarchical genetic structure and high levels of differentiation are known to lead to a high number of false positives in genomic studies of local adaptation, as observed here (Bierne *et al.*, 2013; Hoban *et al.* 2016).

The markers density obtained with RAD-Seq may be also insufficient to detect a RAD-tag in linkage disequilibrium with a selected locus (Lowry *et al.*, 2017; McKinney *et al.*, 2017; Catchen *et al.*, 2017). With a genome size of about 500 Mb for the red coral (Ganot *et al.*, 2016), we expected a SNP sampling of 1 for 18 kb after the SNPs filtering steps (55 SNPs per Mb). Nevertheless, a potential high linkage disequilibrium may be present in here because of drift, and we thus expected that our RAD-tags to at least detect a signal of genetic adaptation, which remains to be confirmed. Approaches dedicated to the study of polygenic adaptation (Daub *et al.*, 2013) or to the genomic distribution of F_ST_ or nucleotide diversity (Hohenlohe *et al.*, 2010) could be interesting here used, but a reference genome is still lacking for the red coral. On a more theoretical point of view, outlier loci could be linked to intrinsic genetic incompatibilities whose allelic frequencies coupled with environmental barriers (Bierne *et al*., 2011). The frequency of genetic incompatiblities in marine populations is largely unknown but probably under-estimated (Plough *et al.*, 2016). Even if not directly linked to local adaptation, such loci are important factors in the evolution of red coral populations.

### Local adaptation to depth in the red coral

We focused on candidate loci meeting the following criteria: i) detection with ARLEQUIN and pcadapt, ii) significant differentiation between depth, iii) function relevant to the adaptation to thermal regime. These loci are the most relevant as factors of local adaptation.

The absence of candidate SNPs common to the three geographical regions could indicate that the adaptation to comparable shallow environmental pressures in these independent regions are based on different genetic pathways, or on non-genetic mechanisms (Putnam and Gates, 2015). However, most candidate loci should be in linkage disequilibrium with selected loci, and such association can easily be lost between distant locations through recombination. Differences in the strength of selective pressure in the three regions could also explain the differences in the detected loci. In Marseille we evidenced a clear signal of differentiation between depths according to multivariate and outlier loci analyses. This detection of a signal of local adaptation in the Marseille region is consistent with the observations from studies of thermotolerance differences in this region (Torrents *et al.*, 2008; Haguenauer *et al.*, 2013; Ledoux *et al.*, 2015; Pratlong *et al.*, 2015). In the case of the Marseille region there are then strong evidences of the existence of adaptive differentiation at a scale of few tens of meters only. Concerning Corsican populations, Ledoux *et al*. (2015) reported no phenotypic signal of local adaptation after reciprocal transplant experiment. Here, the most promising candidate for the adaptation to thermal regime, was an homologous to an allene oxide synthase-lipoxygenase which is known to be involved in the response to thermal stress in octocorals (Lõhelaid *et al.*, 2015). This indirect argument would support the presence of local adaptation in this area as well, but more experimental analyzes will be necessary to confirm the involvement of this function in adaptation to thermal stress in this species. Finally, the detection of a reduced number of candidate loci (two) in the Banyuls region would be consistent with the weaker selective pressure here (see above, Table 2).

Previously, we have sequenced the transcriptome of individuals from the two depths of the Marseille site studied here in Marseille (Pratlong *et al.*, 2015). Several genes were differentially expressed between individuals from the two depths outside thermal stress conditions. Some of these genes, such as those from the Tumor Necrosis Factors Receptor Associated Factors (TRAF) family, have been identified as involved in the response to thermal stress in the hexacoral *Acropora hyacinthus* (Barshis et al., 2013). However, none of these differentially expressed genes were identified in our RAD-Seq study. Additionally the differences of expression may result from acclimatization, and are not necessarily adaptive. The use of a reference genome would be useful here as well to study the potential link between candidate SNPs and genes location and function (Manel *et al.*, 2016).

## CONCLUSION

To our knowledge, the red coral presents among the highest levels of differentiation among studies of local adaptation thought genome scans approaches in marine environment (Bradbury *et al.*, 2010; Limborg *et al.*, 2012; Wang *et al.*, 2013; Chu *et al.*, 2014; Milano *et al.*, 2014; Bay and Palumbi, 2014; Bernardi *et al.*, 2016; Araneda *et al.*, 2016; Guo *et al.*, 2016; Benestan *et al.*, 2016; Bongaerts *et al.*, 2017). This study enabled us to empirically emphasize the limitations in the detection and the interpretation of signals of local adaptation using usual statistical methods in this strongly structured species. Both neutral an adaptive divergence highlighted here demonstrate the genetically singularity of shallow populations of the red coral, especially in the Marseille region were the shallowest populations of this species are found. Together, the strong genetic structure we observed between shallow populations, the low dispersal abilities of the red coral and the local adaptation of these individuals to the highly variable thermal conditions they experiment, raise strong concerns about the evolution of shallow populations and the possibility of loss of adaptive variations in case of mortality events. Extending the genomic study initiated here would be useful to study the evolution of this species in heterogeneous and changing environments. Whatever their origin (genetic or environmental), the different thermotolerance levels observed between depths and populations in the red coral should also be taken into account in future studies of adaptive evolution in this species. These results would also be useful for the management of this harvested species which is present in two international conservation conventions.

## DATA ACCESSIBILITY

The raw DNA sequences are available in the Short Read Archive (SRA) database under accession no. SRR5186771-SRR5187129. The filtered SNP file is available from the Dryad Digital Repository: http://dx.doi.org/10.5061/dryad.rs7bm.

## ACKNOWLEDGEMENTS

This work is a contribution to the Labex OT-Med (n° ANR-11-LABX-0061) funded by the French Government “Investissements d’Avenir” program of the French National Research Agency (ANR) through the A*MIDEX project (n° ANR-11-IDEX-0001-02). This project has been funded by the ADACNI program of the French National Research Agency (ANR) (project n°ANR-12-ADAP-0016; http://adacni.imbe.fr). We thank the ECCOREV Research Federation (FR 3098) for the financial support of part of this study. The project leading to this publication has received funding from European FEDER Fund under project 1166-39417. We thank Nicolas Fernandez and Béatrice Loriod from the Marseille TGML platform for their invaluable help and advice with the preparation of the RAD libraries; the team of the MGX platform for the sequencing of the RAD libraries. The authors thank the UMR 8199 LIGAN-PM Genomics platform (Lille, France, especially Véronique Dhennin) which belongs to the ‘Federation de Recherche’ 3508 Labex EGID (European Genomics Institute for Diabetes; ANR-10-LABX-46) and was supported by the ANR Equipex 2010 session (ANR-10-EQPX-07-01; ‘LIGAN-PM’). The LIGAN-PM Genomics platform (Lille, France) is also supported by the FEDER and the Region Nord-Pas-de-Calais-Picardie. We thank the molecular biology service of the IMBE, the informatic service of the Pytheas Institute, and Frédéric Zuberer of the Pytheas Institute for his support in sampling. We thank the Scandola Natural Reserve, especially Jean-Marie Dominici. We also thank Manuela Carenzi, Pierre-Alexandre Gagnaire, François Bonhomme and Nicolas Bierne for stimulating discussions. Three reviewers, Guillaume Achaz, Lucas Gonçalves da Silva, and an anonymous one, greatly helped us to improve this article. This preprint has been reviewed and recommended by Peer Community In Evolutionary Biology (https://dx.doi.org/10.24072/pci.evolbiol.100061).

---

**Figure S1.**
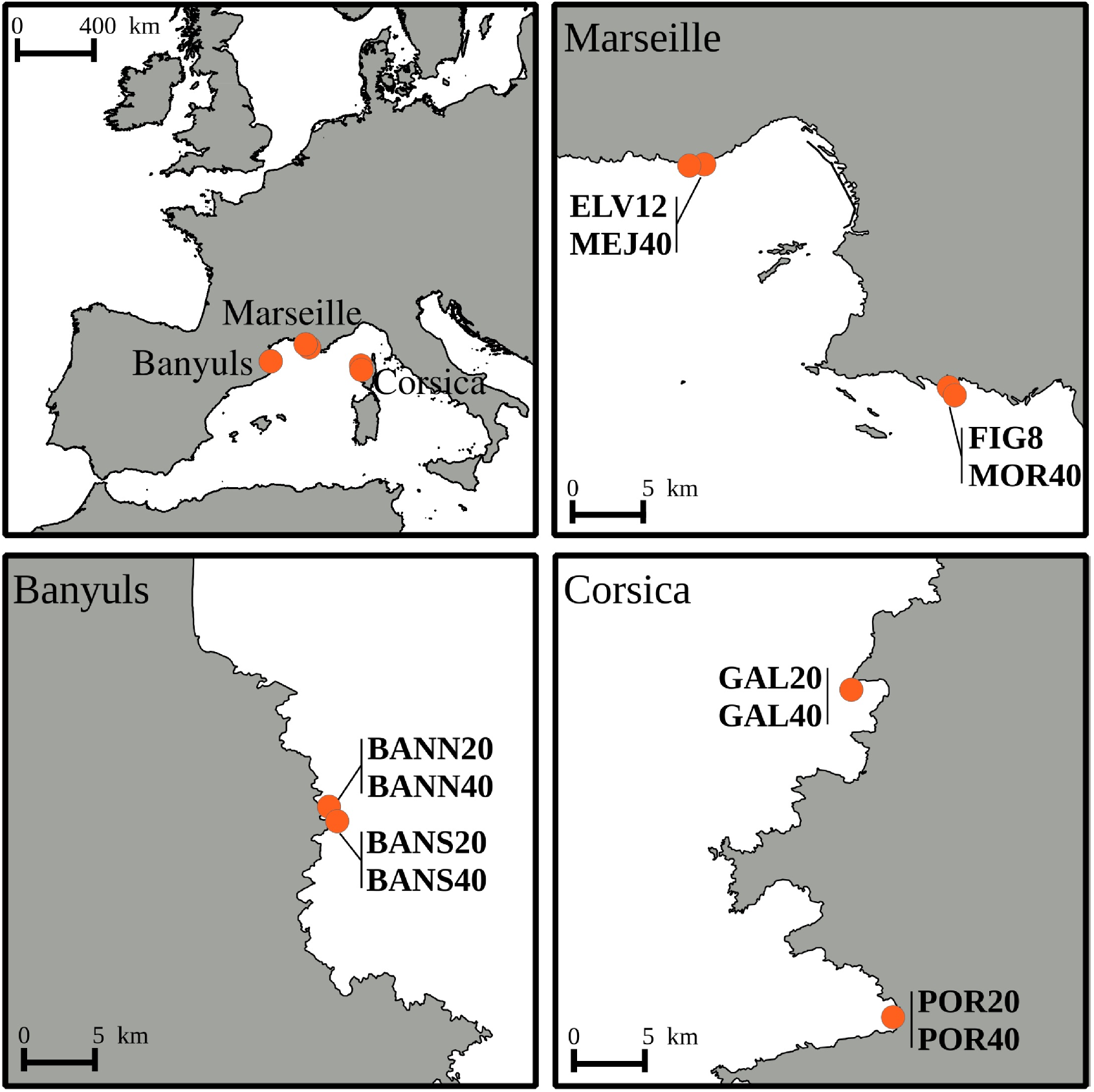
Location of the sampling sites of the red coral among the three studied geographical regions.

**Figure S2.**
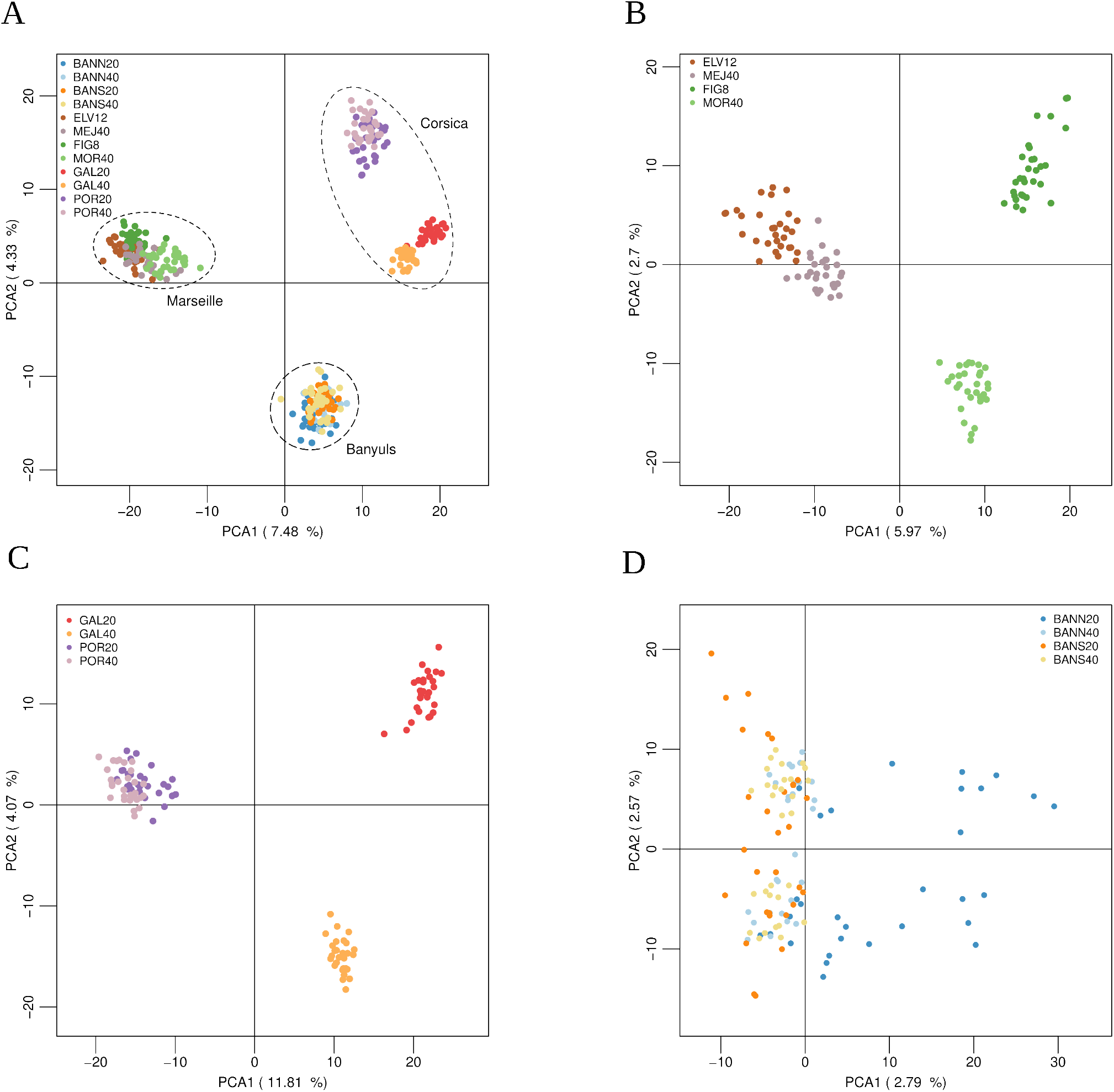
Principal component analysis (Axes 1 and 2), using only putative neutral SNPs, of the A) 12 red coral populations (n = 354 individuals, 25 669 SNPs), B) four red coral populations from Marseille (n =119 individuals, 26 898 SNPs), C) four red coral populations from Corsica (n =117 individuals, 26 592 SNPs), D) four red coral populations from Banyuls (n =118 individuals, 27 069 SNPs).

**Figure S3.**
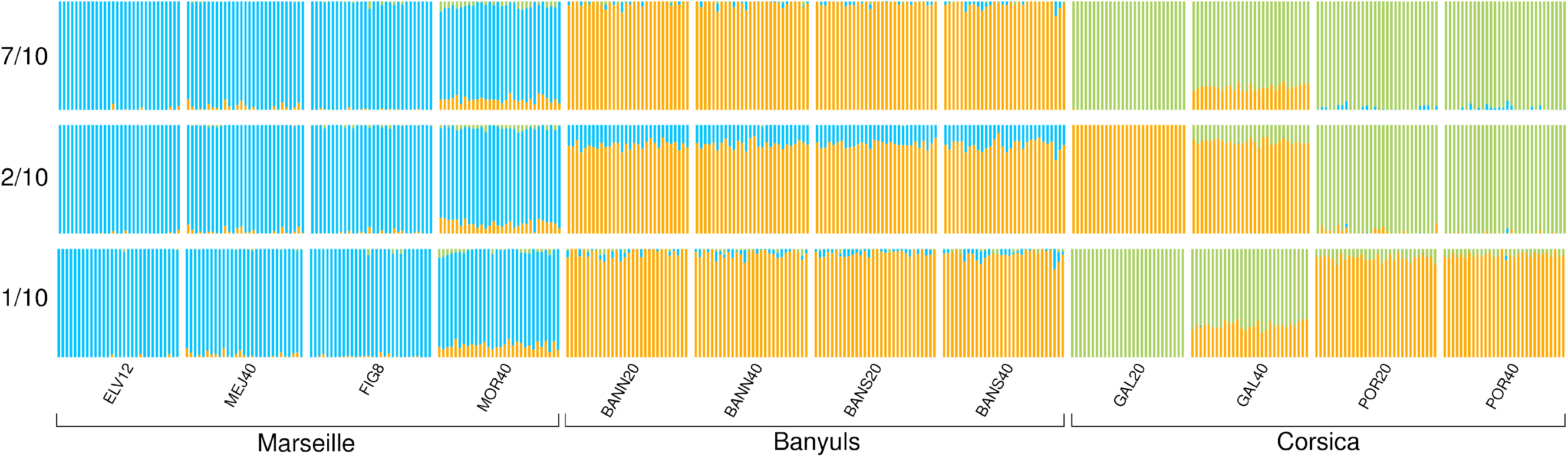
Results from Bayesian individual clustering with STRUCTURE for K = 3. The three figures correspond to major and minor modes detected.

**Figure S4.**
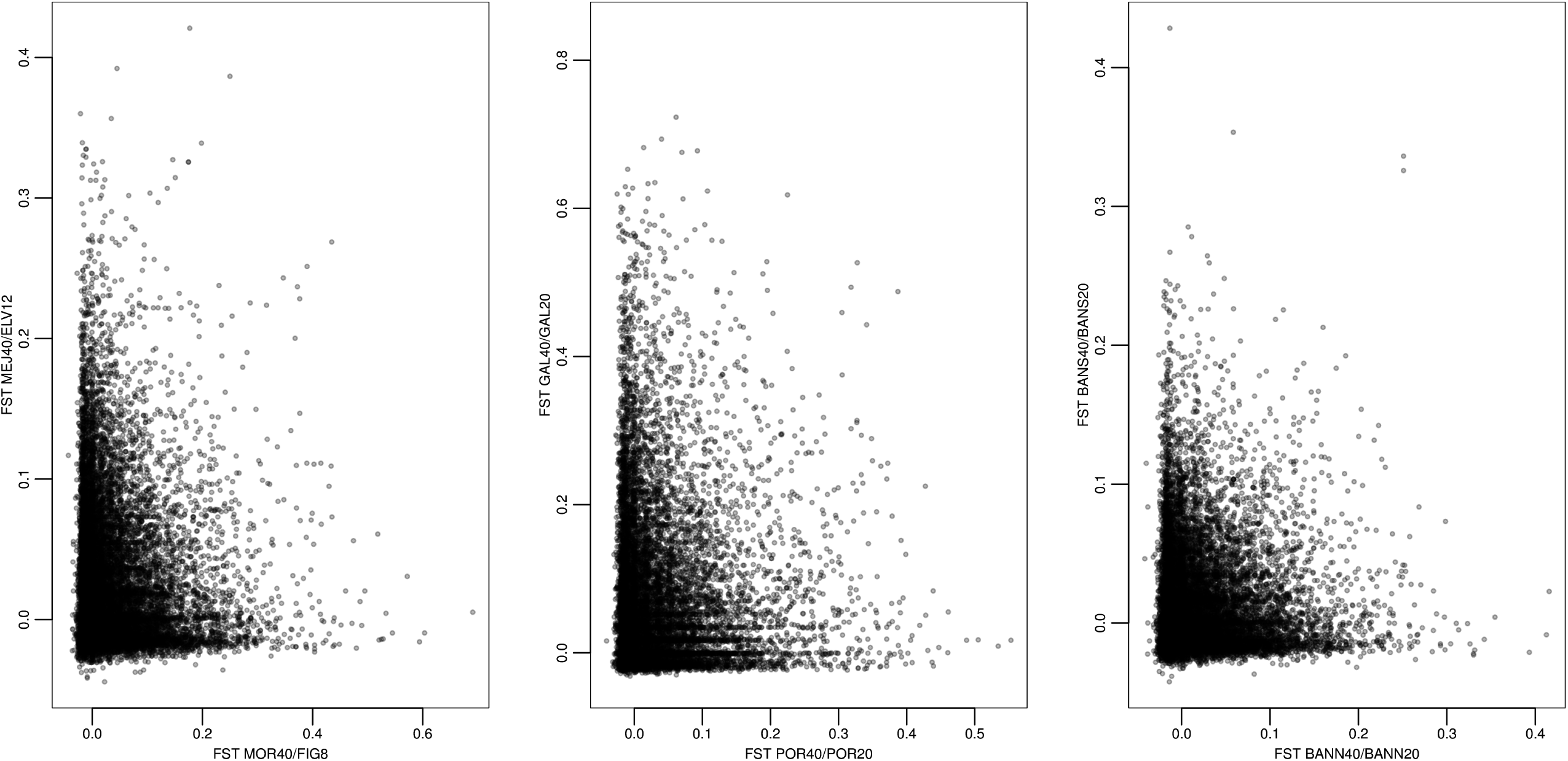
Joint distribution of between-depths F_ST_ in the three geographical regions.

**Table S1.**
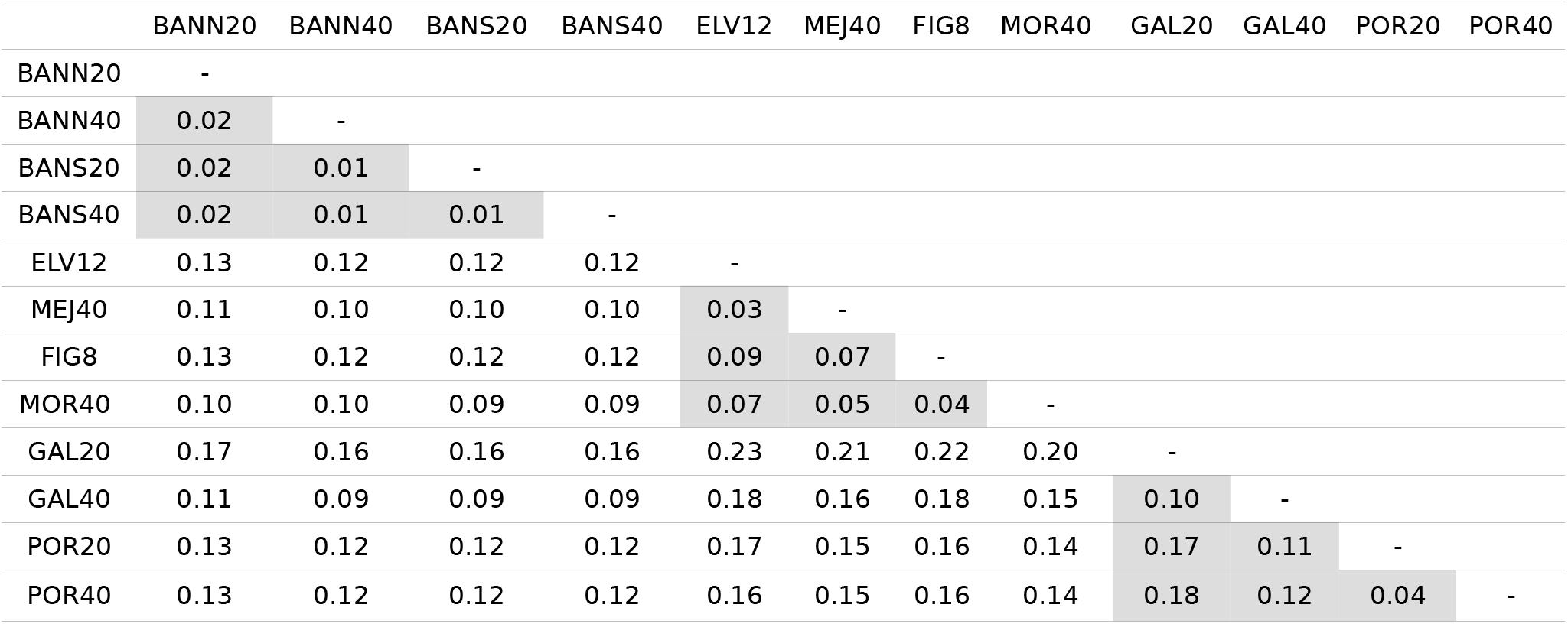
Pairwise F_ST_ estimates using only putatively neutral SNPs. All comparisons were highly significants. Intra-region comparisons are highlighted.

**Table S2.** Pairwise F_ST_ estimates using only outlier SNPs from the ARLEQUIN analysis. All comparisons were highly significants. Intra-region comparisons are highlighted.

|  | BANN20 | BANN40 | BANS20 | BANS40 | ELV12 | MEJ40 | FIG8 | MOR40 | GAL20 | GAL40 | POR20 | POR40 |
| --- | --- | --- | --- | --- | --- | --- | --- | --- | --- | --- | --- | --- |
| BANN20 | - |  |  |  |  |  |  |  |  |  |  |  |
| BANN40 | 0.04 | - |  |  |  |  |  |  |  |  |  |  |
| BANS20 | 0.06 | 0.03 | - |  |  |  |  |  |  |  |  |  |
| BANS40 | 0.06 | 0.03 | 0.02 | - |  |  |  |  |  |  |  |  |
| ELV12 | 0.19 | 0.18 | 0.19 | 0.18 | - |  |  |  |  |  |  |  |
| MEJ40 | 0.16 | 0.14 | 0.15 | 0.15 | 0.04 | - |  |  |  |  |  |  |
| FIG8 | 0.19 | 0.17 | 0.18 | 0.18 | 0.23 | 0.19 | - |  |  |  |  |  |
| MOR40 | 0.14 | 0.12 | 0.13 | 0.13 | 0.19 | 0.14 | 0.08 | - |  |  |  |  |
| GAL20 | 0.27 | 0.27 | 0.28 | 0.27 | 0.37 | 0.35 | 0.36 | 0.32 | - |  |  |  |
| GAL40 | 0.15 | 0.13 | 0.15 | 0.13 | 0.26 | 0.24 | 0.26 | 0.22 | 0.17 | - |  |  |
| POR20 | 0.23 | 0.21 | 0.21 | 0.21 | 0.26 | 0.22 | 0.22 | 0.18 | 0.41 | 0.30 | - |  |
| POR40 | 0.24 | 0.23 | 0.24 | 0.23 | 0.26 | 0.23 | 0.23 | 0.19 | 0.42 | 0.31 | 0.07 | - |

**Table S3.**
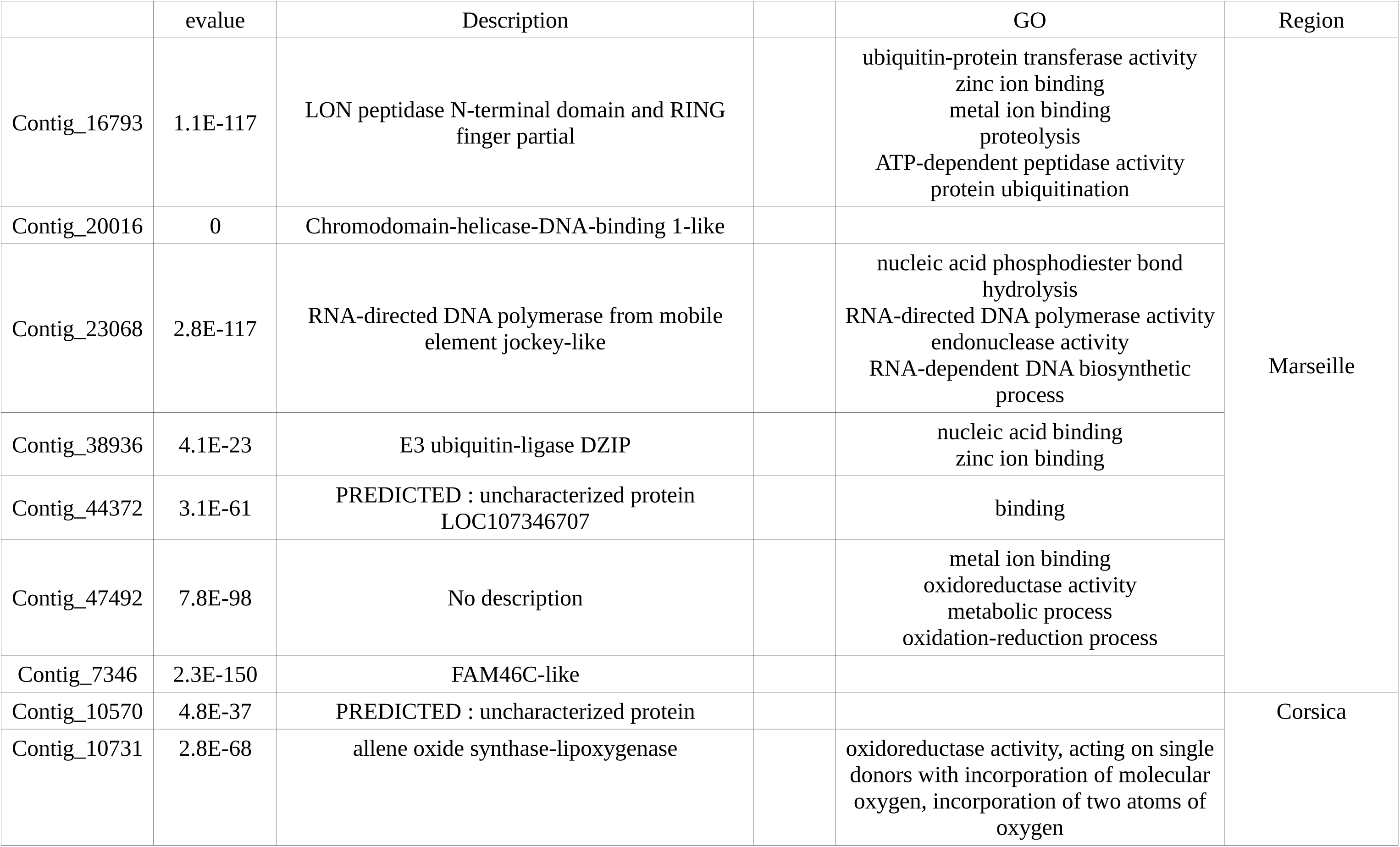
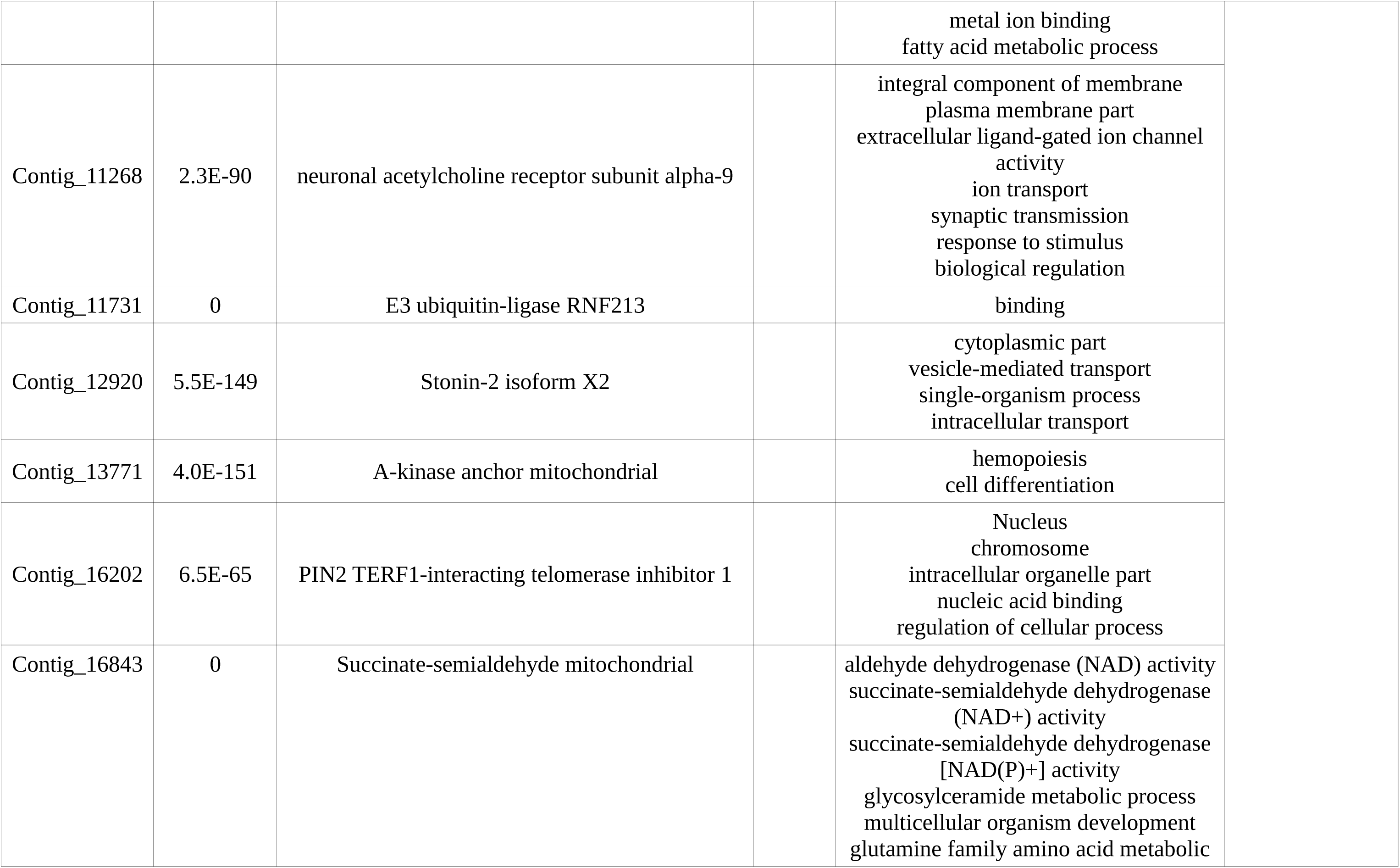
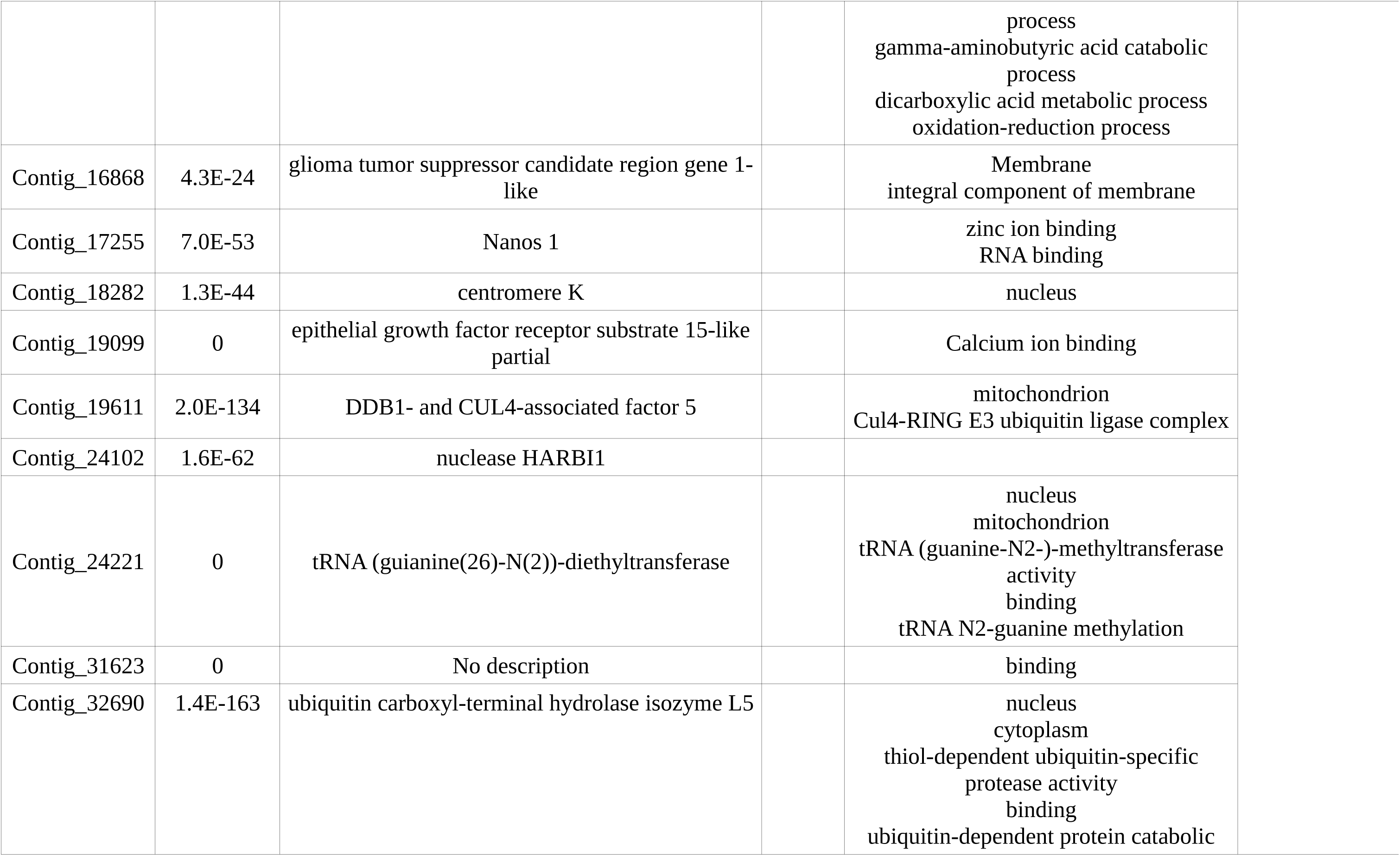
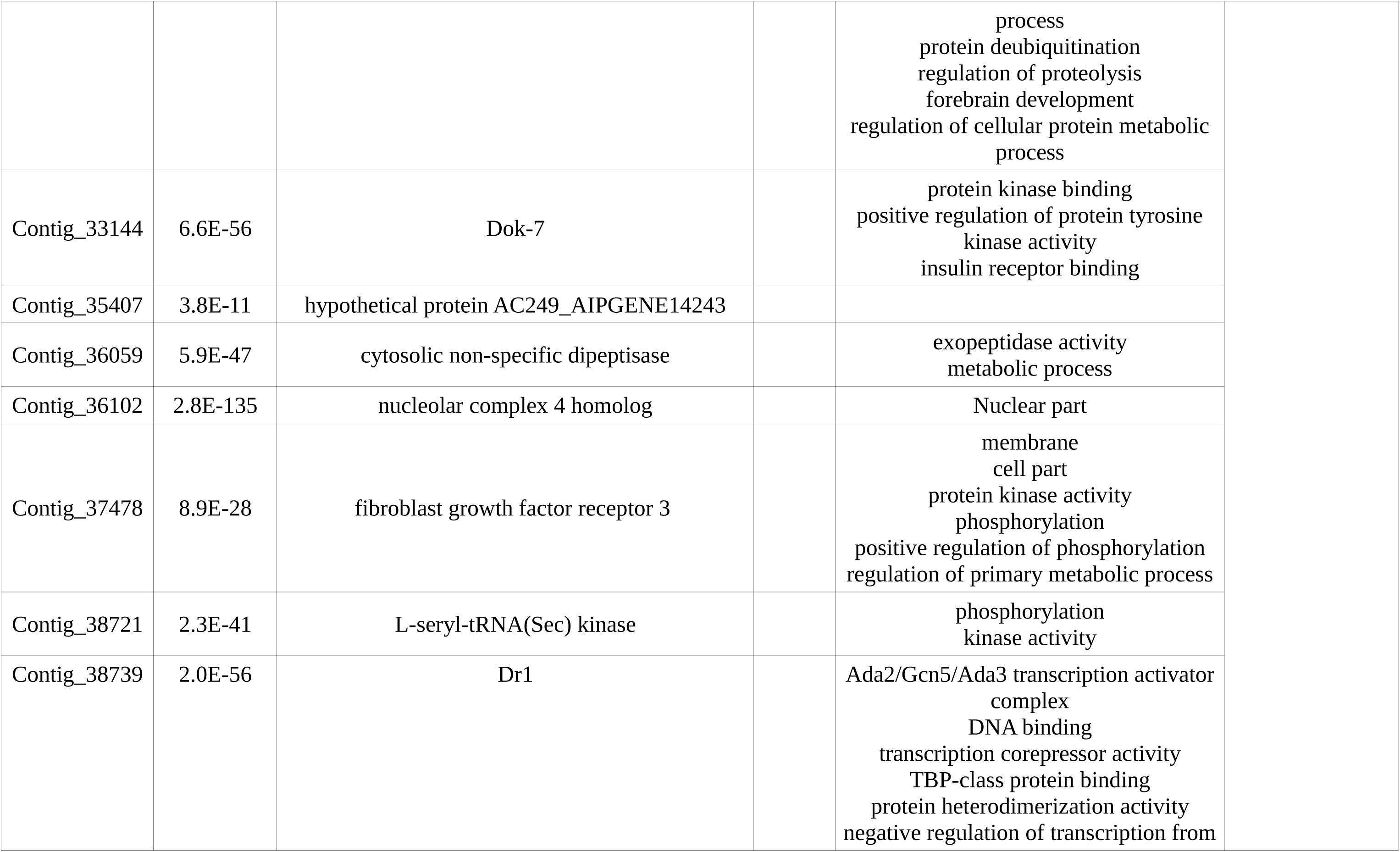
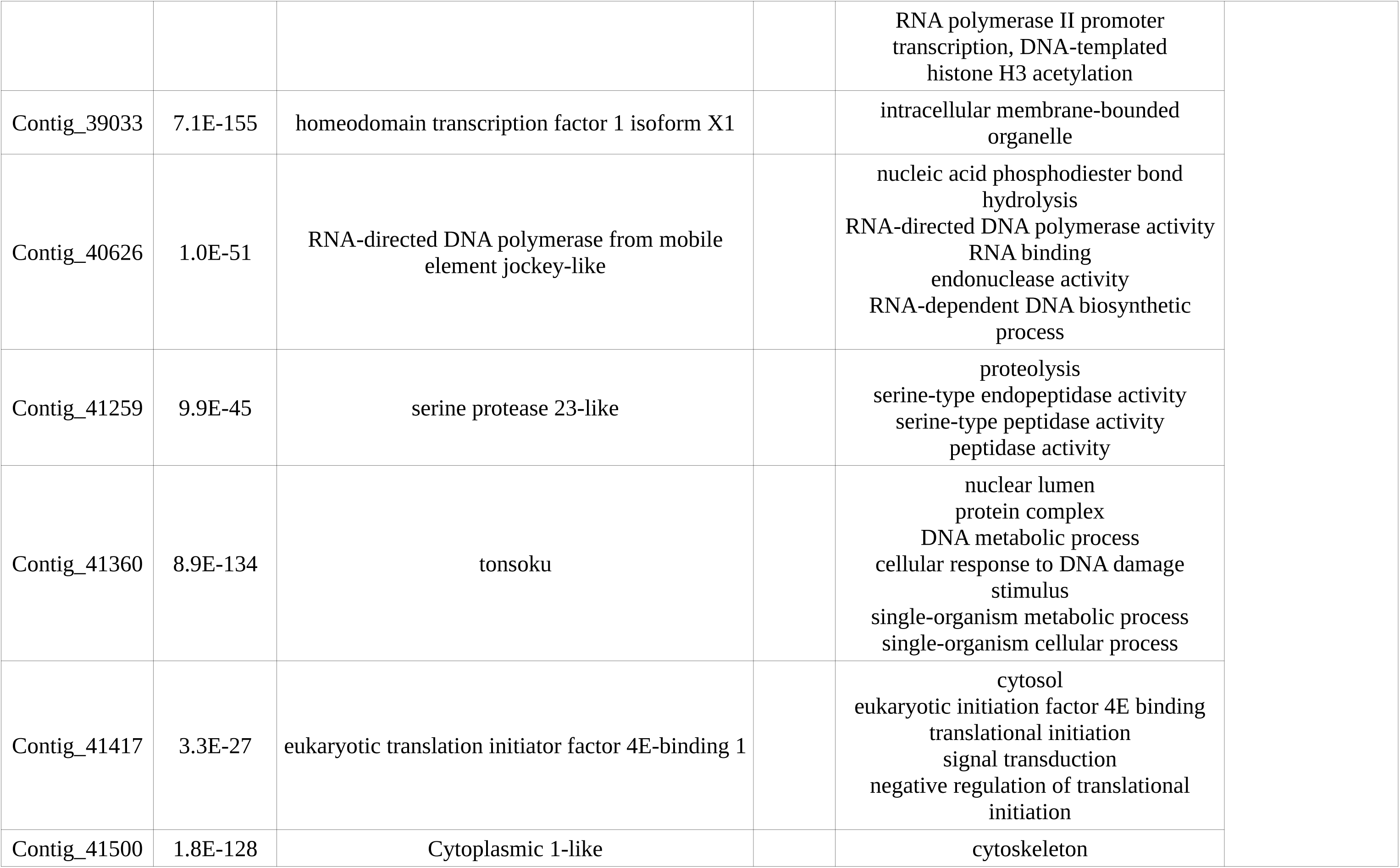
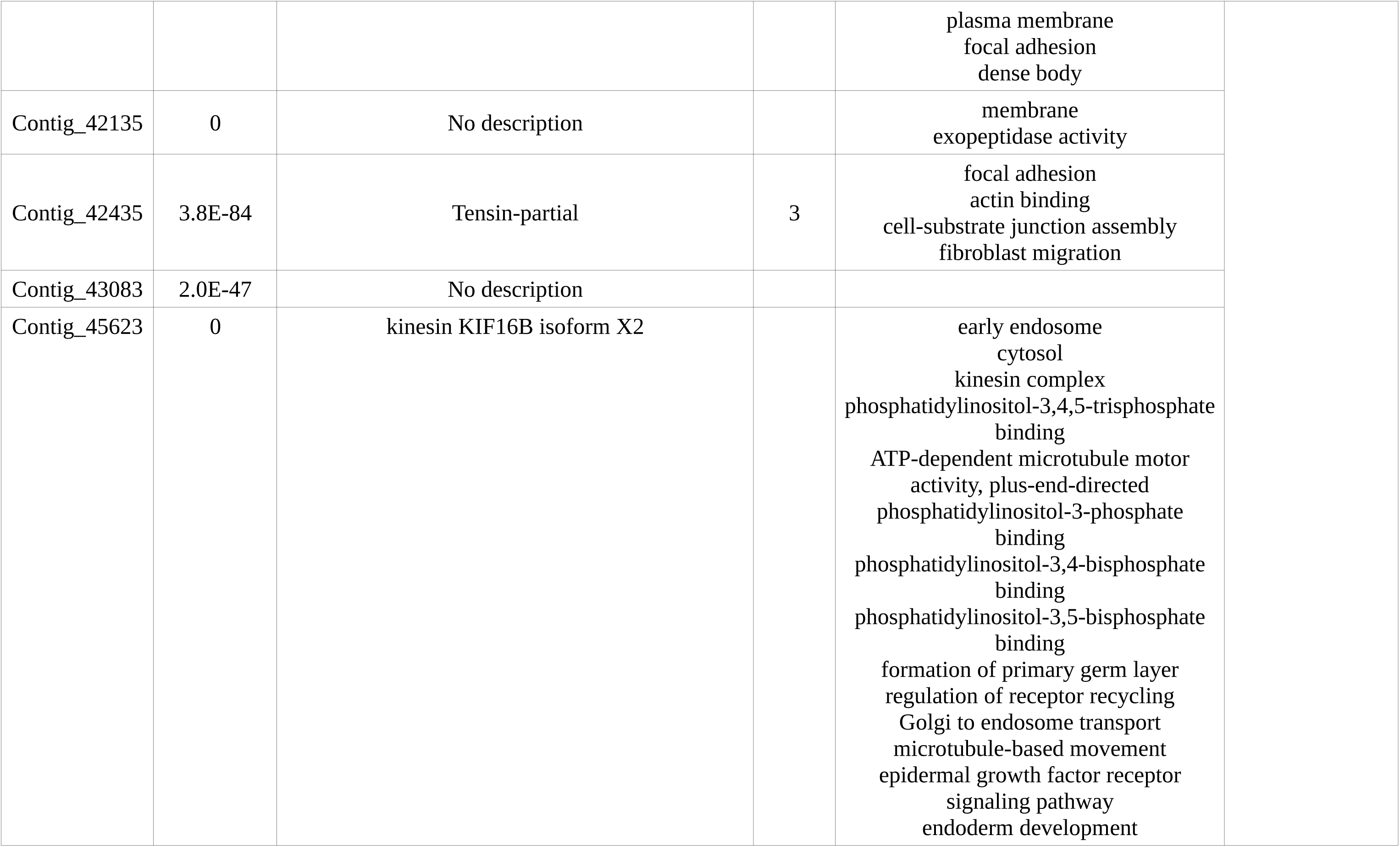
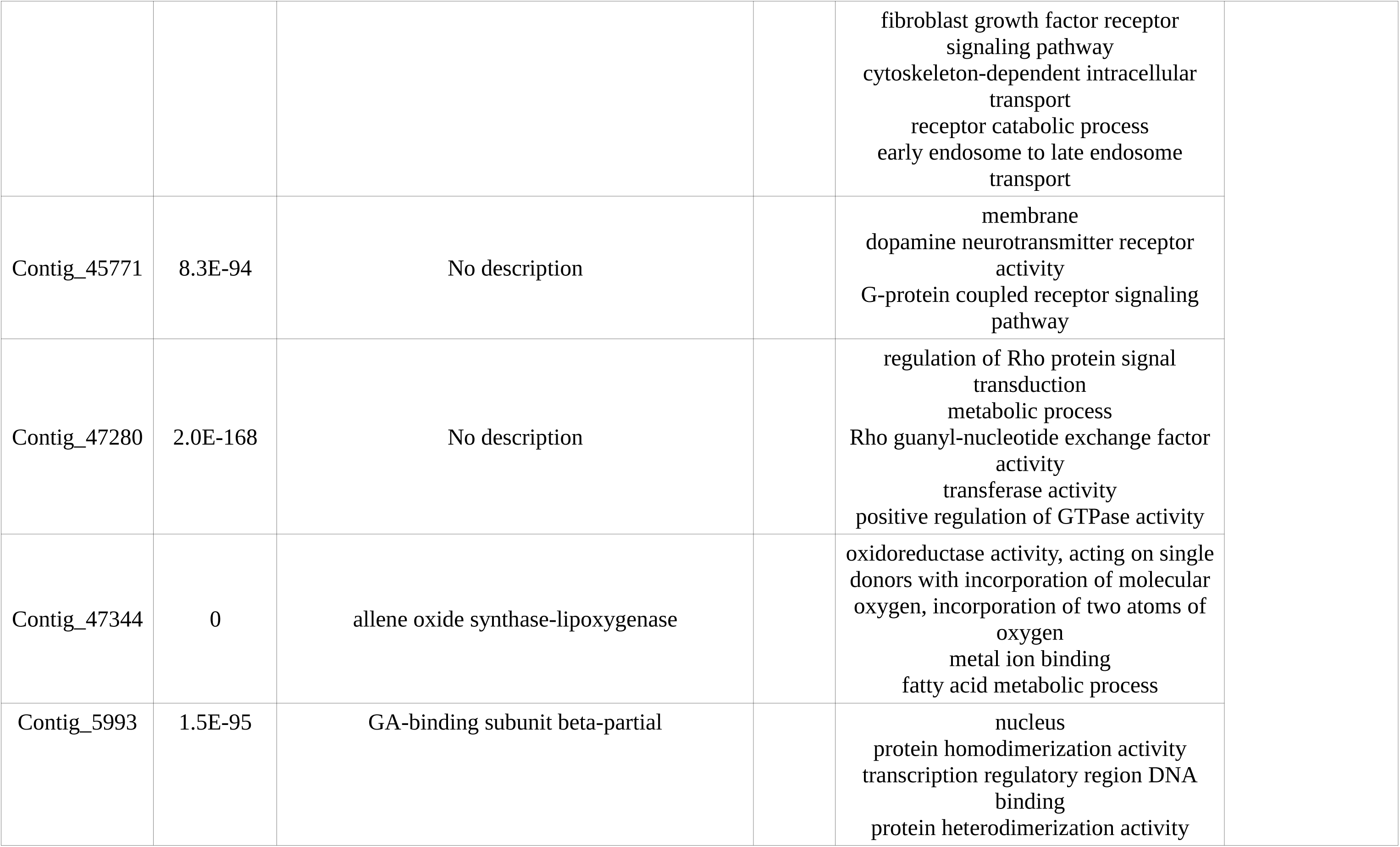
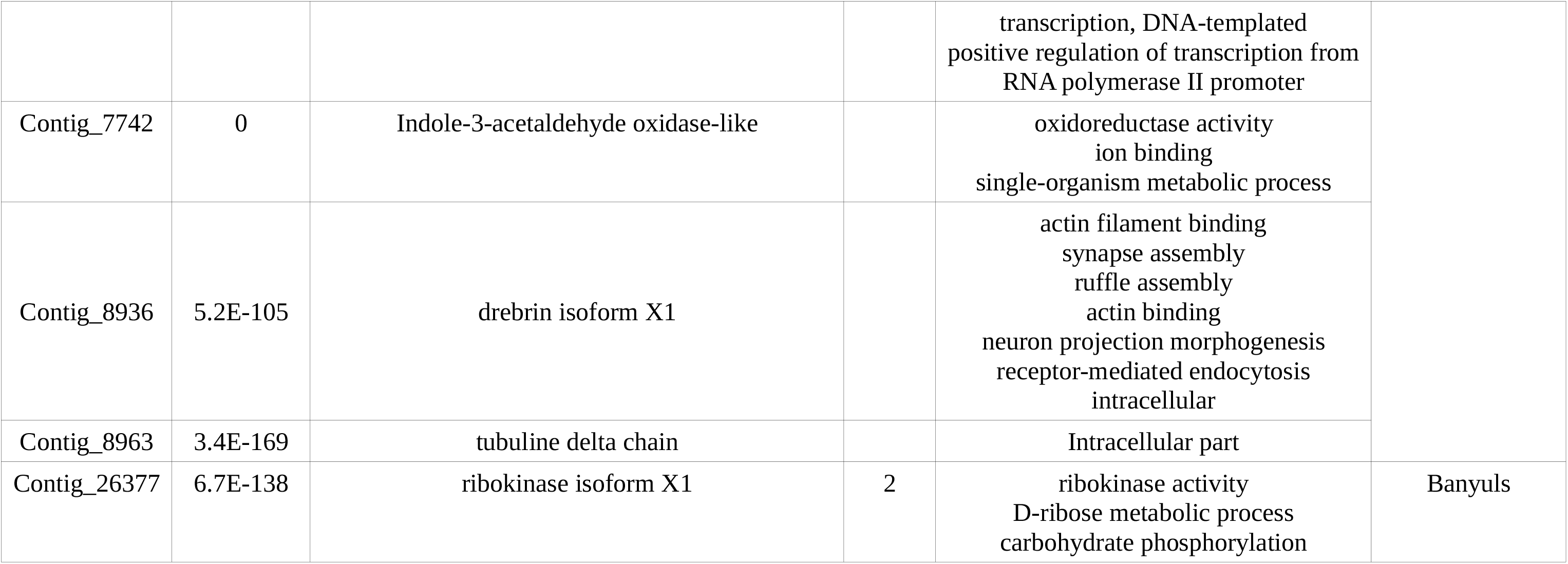
Results of the annotation analysis of candidates for local adaptation in the three geographical regions.

